# Endothelial *GATA6* deficiency suppresses intracellular TLR3-interferon signaling in HPAECs and promotes interferon response in HPASMCs

**DOI:** 10.64898/2025.12.09.693063

**Authors:** Vrinda Dambal, Iram Zaia, Dmitry Goncharov, Tapan Dey, Iryna Zhyvylo, Elena A. Goncharova, Maria Trojanowska

## Abstract

*GATA6* is a key transcription factor crucial for maintaining endothelial cell (EC) homeostasis. The dysregulation of endothelial immune function is a central feature in diseases such as pulmonary arterial hypertension (PAH). In this study, we explored the consequences of *GATA6* deficiency in human pulmonary arterial endothelial cells (HPAECs) and its impact on immune response pathways. We report that siRNA-induced *GATA6* deficiency or the GATA inhibitor led to significant downregulation of interferon response genes and a marked reduction in toll-like receptor 3 (*TLR3*) expression. *GATA6* overexpression enhanced the expression of these genes, and *TLR3* inhibition abrogated this response in HPAECs. Furthermore, conditioned medium (CM) from *GATA6*-deficient HPAECs upregulated interferon genes in pulmonary artery smooth muscle cells (HPASMCs), indicating a paracrine effect. Overall, these findings highlight the critical role of *GATA6* in modulating *TLR3* signaling and immune responses in endothelial cells and suggest its involvement in endothelial-smooth muscle cell interactions in vascular inflammation.

**New and Noteworthy:** We show that endothelial *GATA6* is required for proper activation of intracellular *TLR3*-interferon signaling in HPAECs. *GATA6* loss diminishes interferon pathway responses in endothelial cells while promoting an exaggerated interferon signature in adjacent smooth muscle cells via secreted factors. This work reveals a new GATA6-dependent mechanism governing endothelial–smooth muscle crosstalk in pulmonary vascular disease.

## Introduction

ECs are the innermost layer of blood vessels providing the first line of defense against various exogenous substances in the blood, such as cytokines, bacteria, and dsRNA, which can act as immunological triggers. The ECs in the blood vessels respond to injury by secreting chemokines, cytokines and by expressing adhesion molecules.^1–3^.

ECs play an important role in innate immunity and mediate inflammation through pathogen recognition receptors (PRRs), primarily through Toll-like receptors (TLRs).^4^ *TLR3* is a member of TLR family of receptors that has been implicated in several cardiovascular diseases including PAH. It has been shown that *TLR3* expression is reduced in the lungs and PAECs from PAH patients ^5^. Additionally, the apoptotic ECs in human PAH lesions exhibit a loss of *TLR3* expression. These findings point to a protective role of *TLR3* in pulmonary vasculature.^5^ *TLR3* is activated by dsRNA from viruses or a synthetic dsRNA ligand like polyinosinic-polycytidylic (polyIC). It also recognizes siRNAs (small interfering RNAs) and self-RNAs from damaged cells.^6–9^ Once dsRNA blinds to endosomal *TLR3*, it recruits TRIF (toll-interleukin-1 receptor domain containing adaptor inducing interferon-beta (IFN-B)). These signaling proteins then lead to the activation of transcription factors interferon regulatory factor (*IRF3/7*), nuclear factor kappa-B(*NF-κB*) and activator protein 1 (*AP-1*), which, in turn, mediate the production of type 1 interferons (IFNs), proinflammatory cytokines and chemokines.^10^ Vascular smooth muscle cells (VSMCs) also express *TLR3* abundantly and *TLR3* is found to be upregulated in atheroma-derived SMCs.^11,12^.

*GATA6* belongs to the GATA family of transcription factors that bind to DNA sequence (A/T) GATA(A/G).^13^ *GATA6* was found to be highly expressed in human umbilical vein endothelial cells (HUVECs) where it regulates expression of genes involved in maintaining the vascular tone, such as *NOS3* and *ET-1*. In addition, it also regulates VCAM-1 (Vascular cell adhesion molecule-1), which is expressed in response to inflammation^14–16^. *GATA6* is important for regulation of vessel fate and vascular remodeling and its expression is lost during vascular injury.^17^ Previously, we demonstrated that *GATA6* is reduced in PAECs and PASMCs from patients with idiopathic PAH and scleroderma-PAH. We also found that mice with conditional endothelial GATA6 deletion (GATA6 CKO) spontaneously develop pulmonary vascular remodeling and PH.^18^ In addition, *GATA6* deficiency was found to cause oxidative stress and mitochondrial dysfunction as it suppressed the expression of antioxidant enzymes *SOD2, GPX1, GPX7*, among others, in HPAECs as well as the GATA6 CKO mice. Likewise, when *GATA6* was depleted from healthy PASMCs, there was a significant increase in mitochondrial and cellular reactive oxygen species levels. On the contrary, when *GATA6* was overexpressed in the PASMCs from PAH, there was a significant reduction in proliferation and induction of apoptosis. A bulk RNA seq of PASMCs from patients transduced with *GATA6* adenovirus show an upregulation of genes that inhibit PASMC proliferation and a downregulation of pro-proliferative genes *MYEOV* and *PRPF4*, as well as upregulation of growth suppressor genes *TMEM173* and *EAF1*.^19^

While these previous studies have shown the importance of *GATA6* in regulating endothelial and smooth muscle cell function, its role in modulating immune response in endothelial and smooth muscle cells has not been explored. Our work here shows that *GATA6* deficiency in HPAECs leads to the downregulation of *TLR3*-mediated signaling pathways, impairing the production of interferons and cytokines critical for immune response. Moreover, conditioned medium derived from *GATA6*-deficient HPAECs affects HPASMCs, by activating *TLR3* signaling in these cells. This crosstalk between endothelial and smooth muscle cells could have significant implications for vascular inflammation and pulmonary vascular diseases, including PAH.

## Methods

### Human cell cultures

Primary HPAECs were obtained from two independent vendors to ensure reproducibility across donor backgrounds (Cell applications, Promocell). HPAECs were maintained in complete Endothelial Cell Growth Medium MV2 (PromoCell, C-22221) supplemented with the corresponding supplement mix (PromoCell, C-39221). HPASMCs (Cell Applications Inc.) were isolated from the same donor as the HPAECs, providing a matched endothelial–smooth-muscle pair for paracrine and coculture experiments. Additional HPASMCs were provided by the University of Pittsburgh Vascular Medicine Institute Cell Processing Core and were maintained in LONZA Smooth Muscle Growth Medium-2 (SMGM-2) supplemented with 100 U/mL penicillin and 0.1 mg/mL streptomycin.^20^ Human lung microvascular endothelial cells (HMVEC-L; Lonza) were used in selected comparative experiments; donor demographics were not disclosed by the vendor. (Table 1). Cell identity was further verified by the expression of endothelial and smooth muscle markers (CD31 and α-SMA, respectively; Supplementary Fig. 5D–E(https://doi.org/10.6084/m9.figshare.30680609.)). All cultures were between passages 4–7. Biological replicates (n ≥ 5) represent independent cultures.

**Table 1:** Human cell lines.

| Cell Type | Vendor / Source | Catalog / Lot Number | Donor Information |
| --- | --- | --- | --- |
| HPAEC<br>(PromoCell) | PromoCell | 458Z016.12 | 51-year-old Caucasian female, main pulmonary artery |
| HPAEC (Cell Applications) | Cell Applications Inc. | Lot 1487 | 21-year-old Caucasian male, motor-vehicle accident, marijuana-positive toxicology |
| HPASMC<br>(Cell Applications) | Cell Applications Inc. | Lot 1487 | Matched donor with Cell Applications HPAECs (Lot 1487) |
| HPASMCs | University of Pittsburgh Vascular Medicine Institute Cell Processing Core | M | 38 year old Male |
| HMVEC-L<br>(Lonza) | Lonza | N/A | N/A |

### RNA isolation and quantitative PCR

Trizol (15596026, Fisher) was used to isolate RNA from HPAECs and mouse endothelial cells. RNA was isolated using the Zymo Quick-RNA Miniprep Kit (R1055, Zymo), following manufacturer’s recommendation. RNA was quantified using the nano-spectrophotometer; 1μg of total RNA was used to reverse transcribe the RNA into cDNA using the High-Capacity cDNA Reverse Transcription kit (4374966, Applied biosystems). The diluted cDNA was used to measure gene expression quantitatively using the SYBR Green PCR kit (4309155, Fisher) and the Step OnePlus Real-Time PCR system. Relative gene expression of genes of interest was measured through the 2−ΔΔCT method amongst the treatment and control groups and normalized to the housekeeping genes (*GAPDH*). Primers are listed in Table 2.

**Table 2:**
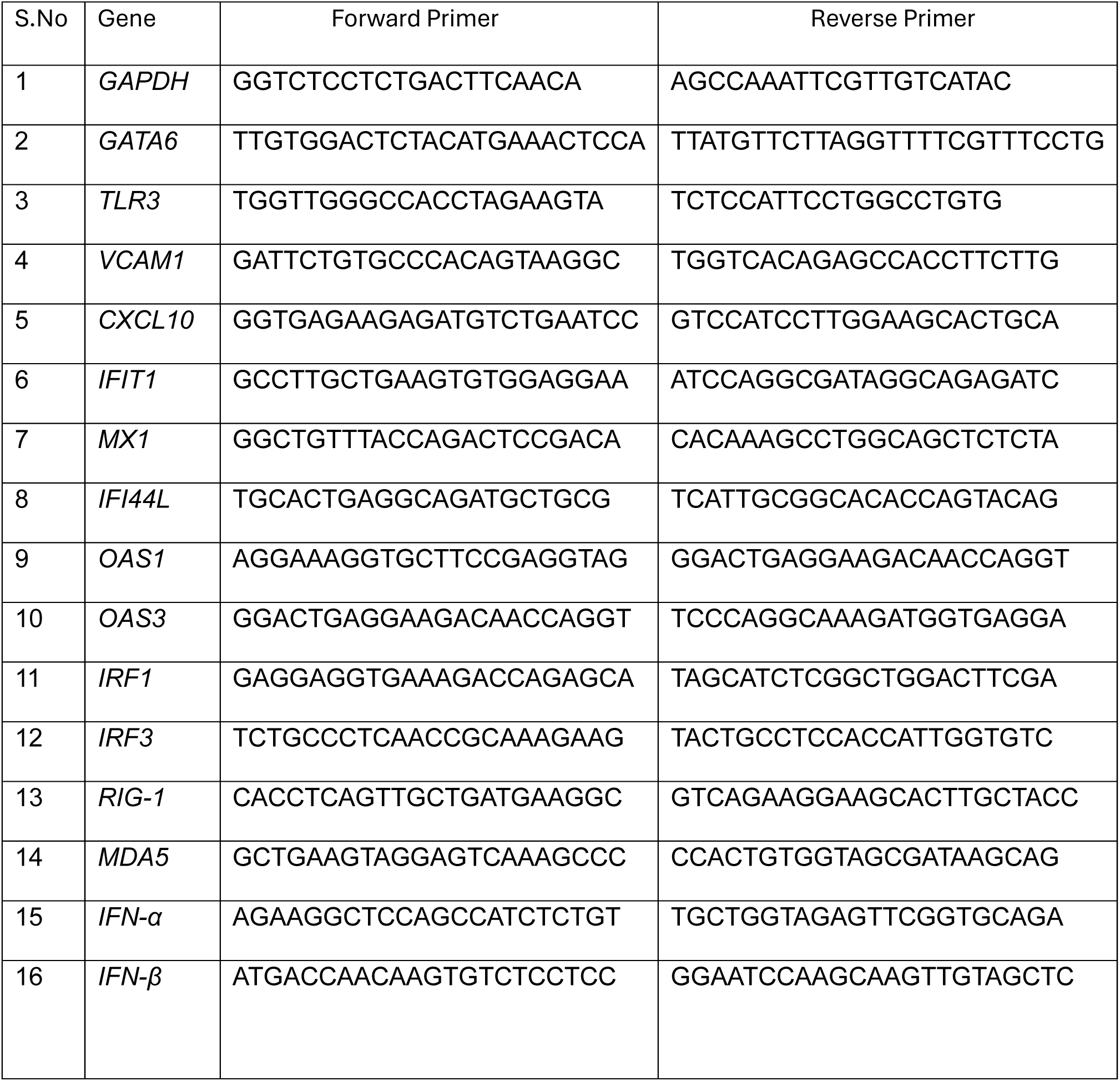
Human qPCR Primers.

### Microarray analysis

Microarray analysis was performed at The Boston University Microarray Core Facility. Affymetrix CEL files were normalized to produce gene-level expression values using the implementation of the Robust Multiarray Average (RMA) in the affy package (version 1.36.1) included within in the Bioconductor software suite (version 2.12) and an Entrez Gene-specific probe set mapping from BrainArray (version 16.0.0). RLE and NUSE quality metrics were computed using the affyPLM Bioconductor package (version 1.34.0). All microarray analyses were performed using the R environment for statistical computing (version 2.15.1). (PMID:37087509)

### RNA sequencing

RNA sequencing (RNA-seq) was performed at the Boston University Microarray and Sequencing Resource core facility. FASTQ files were aligned to human genome build hg38 using STAR (version 2.7.9a)^21^. Ensembl-Gene-level counts for non-mitochondrial genes were generated using featureCounts (Subread package, version 2.0.3) and Ensembl annotation build 112 (uniquely aligned proper pairs, same strand). FASTQ quality was assessed using FastQC (version 0.11.7), and alignment quality was assessed using RSeQC (version 3.0.0). Variance-stabilizing transformation (VST) was accomplished using the variance Stabilizing Transformation function in the DESeq2 R package (version 1.32.0)^22^. Differential expressions were assessed using the Wald test implemented in the DESeq2 R package. Correction for multiple hypothesis testing was accomplished using the Benjamini-Hochberg false discovery rate (FDR). All analyses were performed using the R environment for statistical computing (version 4.1.2).

### Gene Set Enrichment Analysis (GSEA)

GSEA (version 2.2.1)^23^ was used to identify biological terms, pathways and processes that are coordinately up- or down-regulated in the RNA-seq experiment. The Entrez Gene identifiers of all genes in the Ensembl Gene annotation were ranked by the Wald statistic computed between the siScr and siGATA6 groups. This ranked list was then used to perform pre-ranked GSEA analyses (default parameters with random seed 1234) using the Entrez Gene versions of the H (Hallmark), C2 CP (Biocarta, KEGG, PID, Reactome, WikiPathways), C3 (microRNA and transcription factor targets), and C5 (GeneOntology, GO) gene sets obtained from the Molecular Signatures Database (MSigDB), version 2024.1.Hs. The complete set of GSEA results is in Supplementary Table (https://doi.org/10.6084/m9.figshare.30680609.).

### Ingenuity Pathway Analysis (IPA)

Differentially expressed genes (DEGs) from a previously published comparison of HPAECs treated with siGATA6 or siScr [PMID 37087509] were analyzed using Ingenuity Pathway Analysis (IPA; QIAGEN Inc.). The input dataset included official gene symbols, log₂ fold change, and adjusted *p* values (FDR *q* values) for genes with FDR *q* < 0.05. Core analysis was performed using *Homo sapiens* as the species, including both direct and indirect experimentally validated interactions. Canonical pathway enrichment was evaluated using right-tailed Fisher’s exact test, and activation *z*-scores were used to infer pathway activation or inhibition. Pathways with FDR *q* < 0.05 and |*z*| > 2 were considered significantly affected. Visualization of top enriched pathways was generated using the –log₁₀(*p*) values, gene ratio, and number of overlapping genes. Complete IPA outputs are provided in Supplementary Table S1 (https://doi.org/10.6084/m9.figshare.30680609.).

### Adenoviral constructs

Adenoviral vectors encoding human *GATA6* (AdGATA6) and control green fluorescent protein (AdGFP) were obtained from Vector Laboratories and have been previously described.^24^Cells were transduced with AdGATA6 or AdGFP in serum-free medium for 6 hours, followed by replacement with complete growth medium. Viral dose titration was empirically determined by quantifying adenoviral Ad5 gene expression via qPCR to ensure efficient transduction without cytotoxicity.

### siRNA transfection

HPAECs at ∼80% confluence were transfected with 10 nM small interfering RNA targeting human GATA6 (siGATA6; Dharmacon, L-008351-00-0005) or non-targeting scrambled control siRNA (siScr; Dharmacon, D-001810-01-20) using Lipofectamine RNAiMAX (Thermo Fisher Scientific, 13-778-150) in complete Endothelial Growth Medium MV2 (PromoCell, C-22221). After 24 h, cells were harvested for RNA extraction using TRIzol reagent or lysed in RIPA buffer for protein analysis. Knockdown efficiency was verified by qRT-PCR and Western blotting for GATA6.

### Western Blot

The cells were washed with ice-cold PBS and then collected in a RIPA buffer (89901, Fisher) with 1X protease and phosphatase inhibitor. The protein was quantified using the BCA Assay (23227, Fisher). Equal amounts of protein were loaded on a precast polyacrylamide gel (4561084, BioRad) and transferred on a PVDF membrane (1704272, Fisher). The membranes were blocked with 5% nonfat dry milk in TBST and incubated overnight with the following primary antibodies: goat anti-human *GATA6*(R&D, AF1700, 1:500), *TLR3* rat anti-human (MA5-16273,1:500, Fisher), mouse anti-human *ꞵ-actin* (A1978,1:2000, Sigma). The secondary antibodies used were anti-goat IgG (R&D, HAF109,1:10000), anti-ratIgG (Cytiva,NA935V,1:5000), anti-mouse igG (Invitrogen,62-6520,1:10000). The next day, the membranes were washed with 1X TBST and incubated with HRP-conjugated secondary antibodies at room temperature for one hour. The signal was developed using the West Femto Chemiluminescence kit (34096, Thermofisher). The membranes were reprobed after stripping with the Stripping buffer (21059, Fisher). The band intensity was analyzed by densitometric analysis using ImageJ. The *GATA6* antibody has been validated by our previous studies. Efficient knockdown of *TLR3* was confirmed at protein levels (Supplementary Fig. 5A–B), and antibody specificity was validated using full-length immunoblots with molecular weight markers (Supplementary Fig. 5C) (https://doi.org/10.6084/m9.figshare.30680609.).

### GATA inhibitor (Pyrrothiogatain) treatment

Pyrrothiogatain (3-(2,5-dimethyl-1H-pyrrol-1-yl) thiophene-2-carboxylic acid), a pan-GATA transcription factor inhibitor, was purchased from Santa Cruz Biotechnology (Cat. No. sc-352288). ^25^ HPAECs at approximately 80% confluence were treated with 250 µM pyrrothiogatain or vehicle control (0 µM; equivalent DMSO concentration) for 24 hours in complete Endothelial Cell Growth Medium MV2 (PromoCell).

### PolyIC treatment

HPAECs at approximately 80% confluence were pretreated with 250 µM pyrrothiogatain (Santa Cruz Biotechnology, sc-352288) or vehicle (DMSO) for 1 hour in complete Endothelial Cell Growth Medium MV2 (PromoCell). The medium was then replaced with fresh medium containing pyrrothiogatain (maintaining the same concentration) and 10 µg/mL polyinosinic:polycytidylic acid [Poly(I:C)] (R&D Systems, Cat. No. 4287/10) for 6 hours. Cells were harvested immediately after treatment for RNA extraction and protein lysate preparation.

### TLR3/dsRNA Complex Inhibitor treatment

HPAECs at approximately 80% confluence were pretreated with 5 µM TLR3/dsRNA complex inhibitor [(R)-2-(3-chloro-6-fluorobenzo[b]thiophene-2-carboxamido)-3-phenylpropanoic acid] (Sigma-Aldrich, Cat. No. 614310-10MG) for 1 hour in complete Endothelial Cell Growth Medium MV2 (PromoCell).^26^ Following pretreatment, the medium was replaced with fresh medium containing the same concentration of TLR3 inhibitor along with adenoviral GATA6 (AdGATA6) or control GFP (AdGFP) (Vector Labs) for 6 hours. Cells were harvested immediately after this period for RNA extraction and protein lysate preparation to assess the impact of TLR3 inhibition on GATA6-mediated gene expression.

### TLR3 knockdown and GATA6 overexpression experiment

HPAECs were seeded to reach 70–80% confluence at the time of transfection. Cells were transfected with TLR3-specific siRNA (siTLR3) or non-targeting scrambled siRNA (siScr) (Dharmacon, ON-TARGETplus SMARTpool) at a final concentration of 10 nM using Lipofectamine RNAiMAX (Thermo Fisher Scientific) for 24 h, followed by AdGATA6 or AdGFP infection for 24h.

### Conditioned media experiment

HPAECs at ∼80% confluence were transfected with 10 nM siGATA6 or control scrambled siRNA (Dharmacon) using Lipofectamine RNAiMAX (Thermo Fisher Scientific) for 24 hours in complete Endothelial Cell Growth Medium MV2 (PromoCell). After transfection, the medium was replaced with serum-free MV2 supplemented with 0.1% BSA. The conditioned medium (CM) was collected after 24 hours, centrifuged at 1,000 × g for 10 minutes to remove cellular debris, and stored at 4°C for immediate use.

HPASMCs were cultured to ∼80% confluence and growth-arrested for 18 hours in serum-free medium containing 0.1% BSA. The cells were then treated for 24 hours with CM derived from siScr- or siGATA6-transfected HPAECs. After treatment, HPASMCs were harvested for RNA extraction and downstream gene expression analysis by qRT-PCR.

### Conditioned media experiment with RNAse Treatment

HPAECs at ∼80% confluence were transfected with 10 nM siGATA6 or control scrambled siRNA (Dharmacon) using Lipofectamine RNAiMAX (Thermo Fisher Scientific, 13-778-150) for 24 hours in complete Endothelial Cell Growth Medium MV2 (PromoCell, C-22221). Following transfection, the medium was replaced with serum-free MV2 containing 0.1% BSA, and cells were incubated for an additional 24 hours to generate conditioned media (CM). The collected CM was centrifuged at 1,000 × g for 10 minutes to remove cellular debris.To assess the contribution of extracellular RNA to paracrine signaling, aliquots of CM were treated with 10 µg/mL RNase A (Thermo Fisher Scientific, EN0531) at 37°C for 30 minutes, followed by enzyme inactivation at 70°C for 10 minutes. RNase-treated and untreated CM were then applied to HPASMCs. HPASMCs were cultured to ∼80% confluence, growth-arrested for 18 hours in serum-free medium supplemented with 0.1% BSA and subsequently treated with the respective CM preparations for 24 hours. Cells were harvested for RNA extraction.

### Statistical Analysis

Analyses were performed using Prism 10.0.2 (GraphPad). The values are presented as means ± SE. For analysis of two independent samples, the normality was assessed using the Shapiro-Wilke test followed and Q-Q plot and analyzed for significance using the student’s t-test. The analysis for four or more independent groups was done using one-way ANOVA followed by post-hoc Tukey’s multiple comparison test. In all cases, p < 0.05 were considered significant and are abbreviated in the figures as follows: *p < 0.05, **p < 0.01, ***p < 0.001 and ****p < 0.0001.

## Results

### GATA6 deficiency in HPAECs leads to downregulation of interferon target genes and TLR3

In our previous work, we identified GATA6 as an important transcription factor maintaining EC homeostasis and function.^19^ To further investigate the consequences of *GATA6* deficiency in HPAECs, we analyzed our previously published microarray data of HPAECs treated with scrambled and GATA6 siRNAs^9^. The analysis revealed that several interferon response genes, including *CXCL10, MX2, IFIT1, VCAM1*, and *TLR3*, were strongly downregulated with siGATA6 treatment (Table 3). To validate the microarray findings, we transfected HPAECs with scrambled or GATA6 siRNAs and performed qPCR and immunoblot analyses. The qPCR analysis demonstrated a consistent decrease in the mRNA levels of key interferon response genes *VCAM1*, *CXCL10*, and *TLR3*. In addition, transcription factors *IRF1* and *IRF3* were also downregulated (Fig. 1A), confirming transcriptional suppression of key interferon-response genes. Further validation at the protein level showed a corresponding reduction in TLR3 expression in GATA6 siRNA-treated HPAECs (Fig. 1B, C, Supplementary Figure 1D (https://doi.org/10.6084/m9.figshare.30680609.)).

**Figure 1:**
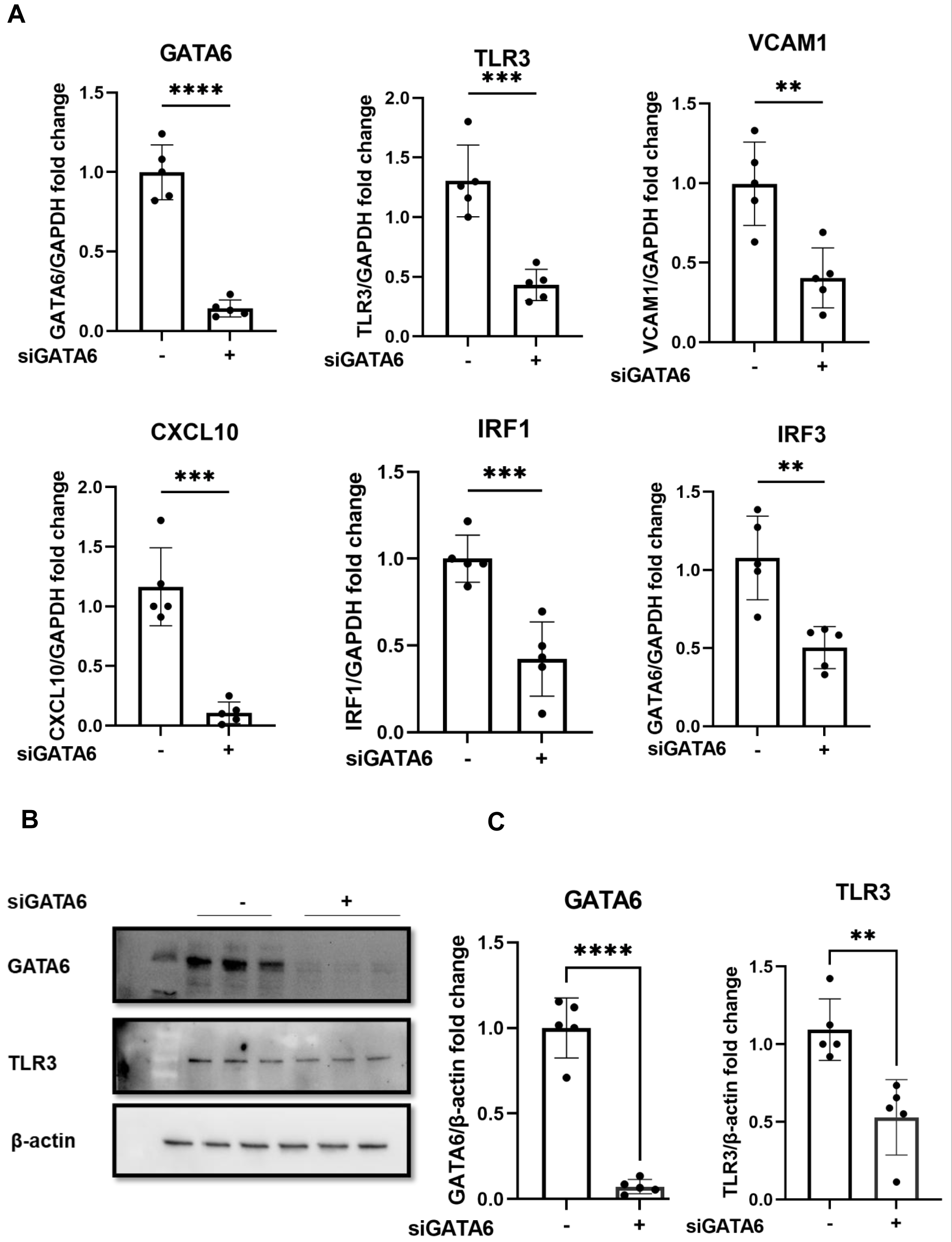
**GATA6 deficiency in HPAECs leads to downregulation of interferon target genes and TLR3.** (A) Quantitative RT-PCR analysis showing mRNA expression of *GATA6, TLR3, VCAM1, CXCL10, IRF1*, and *IRF3* following siRNA-mediated knockdown of GATA6 in HPAECs. Expression levels were normalized to GAPDH and presented as fold change relative to control siRNA-treated cells. Data are means ± SD (n=5, average of 5 independent experiments done in triplicates), Exact p-values: *<u>GATA6</u>* (p < 0.0001), *<u>TLR3</u>* (p = 0.0003), *<u>VCAM1</u>* (p = 0.0034), *<u>CXCL10</u>* (p = 0.0001), *<u>IRF1</u>*(p = 0.0009) and *<u>IRF3</u>*(p = 0.0027) by student t-test. (B) Representative immunoblot analysis of GATA6 and TLR3 protein expression following GATA6 silencing, with β-actin as a loading control. (C) Densitometric quantification of GATA6 and TLR3 protein abundance normalized to β-actin. Data are presented as mean ± SD (n = 5; average of 5 independent experiments). Statistical analysis was performed using unpaired two-tailed Student’s *t*-test; Exact p-values: <u>GATA6</u> (p < 0.0001) and <u>TLR3</u> (p = 0.0106) by student t-test.

**Table 3:** Selected downregulated genes in siGATA6 vs SiScr HPAECs.

| Gene Symbol | siGATA6 vs siScr HPAEC log 2-fold change |
| --- | --- |
| *CXCL10* | -58.22 |
| *MX2* | -38.21 |
| *IFIT1* | -34.95 |
| *RSAD2* | -31.30 |
| *CMPK2* | -30.50 |
| *IFIT3* | -21.35 |
| *CXCL11* | -10.69 |
| *IL-1A* | -8.96 |
| *OASL* | -8.60 |
| *MX1* | -7.45 |
| *IFIT5* | -7.43 |
| *CCL5* | -6.65 |
| *IFIH1* | -6.53 |
| *IFI30* | -6.41 |
| *IFI44L* | -5.62 |
| *IFITM1* | -5.60 |
| *IFI35* | -5.30 |
| *IL8* | -4.43 |
| *VCAM1* | -4.32 |
| *TLR3* | -3.32 |
| *IL6* | -3.26 |

In addition, IPA analysis showed significant inhibition of Toll-like receptor signaling and several pathways related to immune response, including interferon signaling pattern recognition receptors (PRRs) in Bacteria and Virus role of *RIG-1* like receptors in antiviral immunity, in GATA6 siRNA-transfected HPAECs (Supplementary Fig 1A). Taken together, these data indicate that loss of GATA6 in HPAECs down-regulates TLR3 and immune pathways. To test whether this regulatory effect extends across different endothelial subtypes, we repeated *GATA6* inhibition using an additional donor-derived HPAEC line (PromoCell) and human lung microvascular endothelial cells (HMVEC-L; Lonza). The inclusion of HMVECs was intended to address the physiological relevance of distal vascular endothelial cells, which are known to play a major role in pulmonary vascular remodeling and the pathogenesis of PAH. In all cell types, GATA6 depletion led to consistent downregulation of *TLR3*, *VCAM1*, and *CXCL10* transcripts (Supplementary Fig. 1B, C(https://doi.org/10.6084/m9.figshare.30680609.)), indicating that GATA6-dependent modulation of antiviral and inflammatory pathways is a general feature of pulmonary endothelial cells across both conduit and microvascular compartments.

### Pyrrothiogatain treatment downregulates interferon response genes in HPAECs

To further investigate the role of GATA6 in regulating interferon response genes via TLR3 signaling, we examined the effects of pyrrothiogatain, a pan-GATA inhibitor known to block the DNA-binding activity of GATA transcription factors.^25^ Compared with vehicle (DMSO)-treated controls, pyrrothiogatain induced a dose-dependent reduction in the mRNA expression of GATA6-associated targets, including *TLR3*, *VCAM1*, and *CXCL10*, reaching statistical significance at 250 µM (Fig. 2A). This concentration was therefore selected for subsequent experiments.

**Figure 2:**
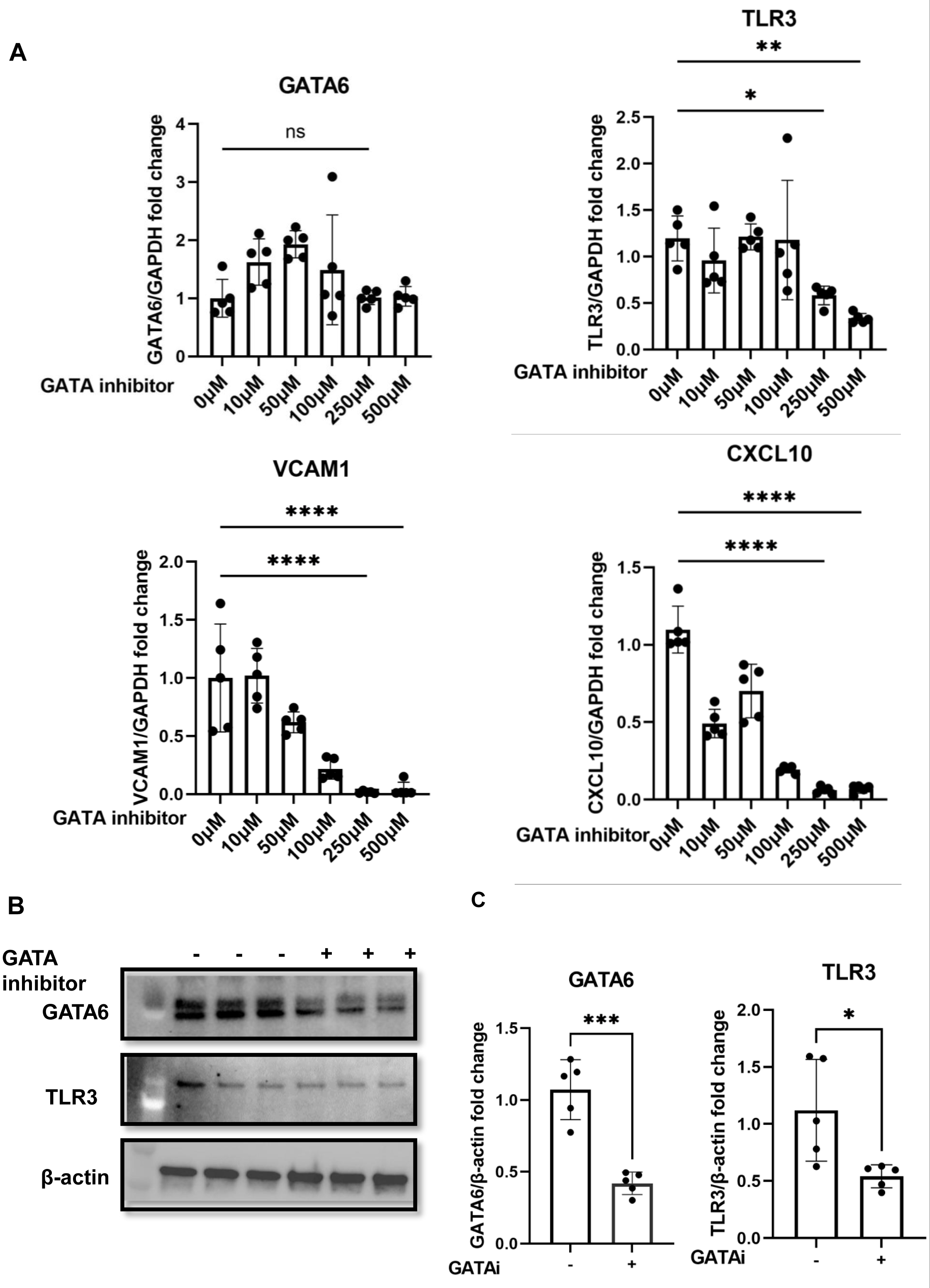
**Pyrrothiogatain treatment downregulates interferon response genes in HPAECs.** (A) mRNA expression levels of *GATA6, TLR3, VCAM1*, and *CXCL10* in HPAECs treated with increasing concentrations of the GATA inhibitor pyrrothiogatain (0–500 µM) for 6hours. Data are presented as mean ± SD (n = 5; three independent experiments performed in triplicate). Exact p-values for each comparison are as follows:*<u>GATA6</u>*: 50 µM (p = ns), 100 µM (p = ns), 250 µM (p = ns), 500 µM (p = ns);*<u>TLR3</u>*: 50 µM (p = ns), 100 µM (p = ns), 250 µM (p = 0.0253), 500 µM (p = 0.0014);*<u>VCAM1</u>*: 50 µM (p = 0.0450), 100 µM (p = 0.0001), 250 µM (p = 0.0001), 500 µM (p = 0.0001); *<u>CXCL10</u>*: 50 µM (p = 0.0001), 100 µM (p = 0.0001), 250 µM (p = 0.0001), 500 µM (p = 0.0001) by one-way ANOVA, Dunnett’s multiple comparisons post-hoc test. (B) Representative Western blot analysis showing protein expression of GATA6 and TLR3 following treatment with vehicle or pyrrothiogatain (250 µM). β-actin served as a loading control. (C) Densitometric quantification of GATA6 and TLR3 protein levels normalized to β-actin. Data are presented as mean ± SD (n = 5; average of 5 independent experiments). Statistical analysis was performed using unpaired two-tailed Student’s *t*-test; Exact p-values: <u>GATA6</u> (p = 0.0002) and <u>TLR3</u> (p = 0.0221) by student t-test.

Consistent with the mRNA data, TLR3 protein levels were significantly reduced following pyrrothiogatain treatment (Fig. 2B, C, Supplementary Figure 2D(https://doi.org/10.6084/m9.figshare.30680609.)). Interestingly, while GATA6 protein abundance was markedly decreased, GATA6 mRNA levels remained unchanged (Fig. 2A, B), suggesting that pyrrothiogatain may additionally impair GATA6 protein synthesis or stability. Together, these findings confirm that pharmacologic inhibition of GATA6 suppresses TLR3 signaling and interferon response genes. Moreover, they indicate that pyrrothiogatain affects GATA6 through dual mechanisms by inhibiting its DNA-binding function and reducing its steady-state protein levels which provides a complementary approach to siRNA-mediated knockdown for dissecting the regulatory role of GATA6 in endothelial immune signaling. These experiments were repeated in an additional donor HPAEC cell line and HMVEC cell line (Supplementary Figure 2A,B(https://doi.org/10.6084/m9.figshare.30680609.)) which exhibited comparable responses. Pyrrothiogatain at 250 µM did not affect HPAEC morphology or confluence, and mRNA levels of apoptosis markers Caspase −3 and Caspase-7 remained unchanged relative to vehicle-treated controls (Supplementary Figure 2C), indicating that the observed effects were not attributable to cytotoxicity.

### GATA inhibition in HPAECs leads to loss of interferon gene response following PolyIC stimulation

Having established a system for GATA inhibition independent of siRNA knockdown, we aimed to further explore the role of *GATA*6 in TLR3 pathway activation. TLR3 signaling was activated using PolyIC, a well-established RNA mimic of viral double-stranded RNA. As anticipated, we found that PolyIC treatment significantly upregulated mRNA levels of *TLR3, VCAM1*, and *CXCL10* in HPAECs. However, GATA inhibitor treatment abrogated these effects, with a down regulation of *TLR3, VCAM1* and *CXCL10* (Fig. 3A). Similar responses were also observed for other RNA-sensing receptors *RIG-1* and *MDA5*, whose PolyIC-mediated induction was suppressed by GATA inhibitor (Fig.3B). This further supports the role of GATA6 in regulating viral sensing pathways. Consistent with the upregulation of the interferon response genes Poly(I:C) treatment robustly induced *IFNα* and *IFNβ* transcripts. However, co-treatment with the GATA inhibitor pyrrothiogatain significantly attenuated this induction, indicating that GATA-dependent transcriptional activity is required for full interferon pathway response (Fig. 3C, D).

**Figure 3:**
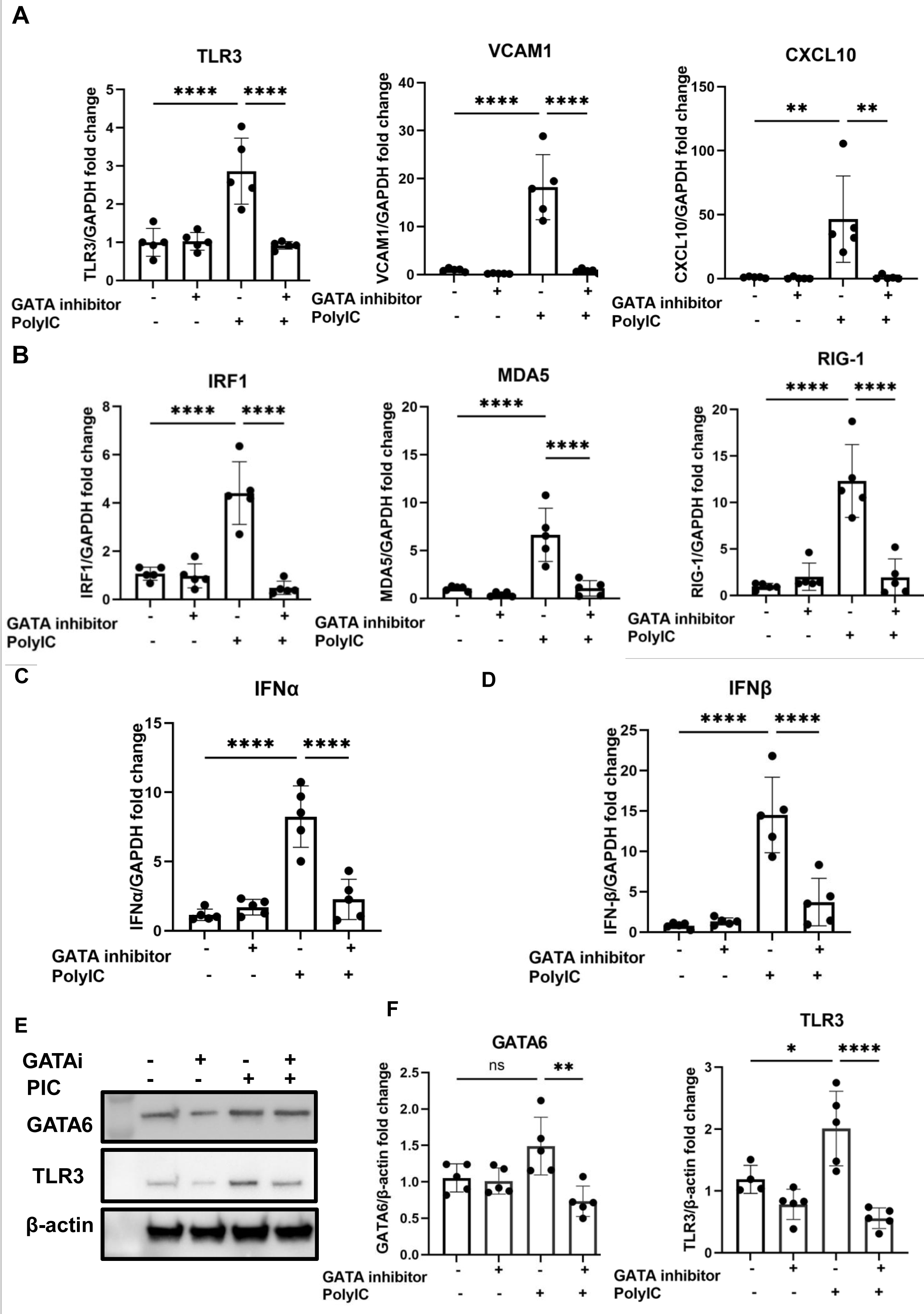
**GATA inhibition in HPAECs leads to loss of interferon gene response following PolyIC stimulation.** (A-D) mRNA levels of *TLR3, VCAM1, CXCL10, IRF1, MDA5, RIG-I, IFNα*, and *IFNβ* were quantified in vehicle-treated and pyrrothiogatain-treated HPAECs with or without PolyIC stimulation (PolyIC ±) using RT-qPCR. Data are presented as mean ± SD (n = 5 independent experiments performed in triplicate). Statistical analysis was conducted using one-way ANOVA followed by Tukey’s multiple comparisons test. Exact p-values were as follows: *<u>TLR3</u>*: vehicle(+)/PolyIC(–) vs vehicle(+)/PolyIC(+), p < 0.0001; vehicle(+)/PolyIC(+) vs inhibitor(+)/PolyIC(+), p < 0.0001. *<u>VCAM1</u>*: p < 0.0001, p < 0.0001. *<u>CXCL10</u>*: p = 0.0030, p = 0.0030. *<u>IRF1</u>*: p < 0.0001, p < 0.0001. *<u>MDA5</u>*: p < 0.0001, p < 0.0001. *<u>RIG-I</u>*: p < 0.0001, p < 0.0001. *<u>IFNα</u>*: p < 0.0001, p < 0.0001. *<u>IFNβ</u>*: p < 0.0001, p < 0.0001. (E) Representative Western blots showing GATA6 and TLR3 protein levels following treatment with GATA6 inhibitor and/or Poly(I:C). β-actin served as a loading control. (F) Densitometric quantification of GATA6 and TLR3 protein abundance normalized to β-actin. Data are presented as mean ± SD (n = 5 independent experiments). p-values were determined using one-way ANOVA with Tukey’s post hoc test. Exact p-values for all pairwise comparisons were as follows: <u>GATA6</u>: vehicle(+)/PolyIC(–) vs vehicle(+)/PolyIC(+), p = ns; vehicle(+)/PolyIC(+) vs inhibitor(+)/PolyIC(+), p = 0.0015. <u>TLR3</u>: p = 0.0015, p < 0.0001.

Additionally, at the protein level, pyrrothiogatain reduced both basal and PolyIC-induced TLR3 protein levels, (Fig 3E, F). Interestingly, Poly(I:C) treatment alone increased GATA6 protein abundance, and this effect was reversed by pyrrothiogatain co-treatment (Fig. 3E, F Supplementary Figure 3C (https://doi.org/10.6084/m9.figshare.30680609.)). This observation suggests a potential feedback mechanism in which TLR3 activation transiently enhances GATA6 expression or stability to reinforce interferon signaling, while pharmacologic GATA inhibition disrupts this positive regulatory loop. To confirm the reproducibility of GATA6-dependent regulation across endothelial subtypes, we repeated the GATA inhibition and PolyIC stimulation experiments in an additional donor-derived HPAEC line and in human lung microvascular endothelial cells (HMVEC-L). HMVECs exhibited comparable responses to those observed in the primary HPAECs, showing robust PolyIC-induced upregulation of *TLR3, VCAM1*, and *CXCL10*, all of which were significantly attenuated by GATA inhibition (Supplementary Fig. 3A) (https://doi.org/10.6084/m9.figshare.30680609.).In contrast, the second HPAEC (PromoCell) line failed to exhibit *TLR3* induction upon PolyIC stimulation and showed blunted regulation of interferon response genes such as *VCAM1* (Supplementary Fig. 3B https://doi.org/10.6084/m9.figshare.30680609). These findings suggest that while the GATA6–TLR3 axis represents a conserved regulatory mechanism in pulmonary endothelial cells, the magnitude of its activation may vary depending on donor-specific factors or endothelial subtype origin.

### Inhibition of TLR3 pathway prevents GATA6-dependent induction of interferon genes

Our study revealed that *GATA6* modulation had a more modest effect on TLR3 expression compared to its pronounced effects on *VCAM1* and *CXCL10* expressions. This observation raised the possibility that GATA6 may regulate these genes directly, independent of TLR3 signaling. To investigate this, we overexpressed GATA6 in HPAECs by infection with GATA6-expressing adenovirus (AdGATA6) in the presence or absence of the TLR3 pathway inhibitor TLR3/dsRNA^26^ and evaluated the expression of interferon response genes. Overexpression of GATA6 significantly increased the expression of *TLR3*, *VCAM1*, and *CXCL10*, whereas the control GFP adenovirus (AdGFP) had a negligible effect. However, the addition of the TLR3 inhibitor completely abrogated the effect of GATA6 overexpression (Fig 4A). These findings suggest that activation of TLR3 signaling is essential for the expression of these genes, and that GATA6 overexpression alone, in the absence of TLR3 activation, is insufficient to induce their expression.

**Figure 4:**
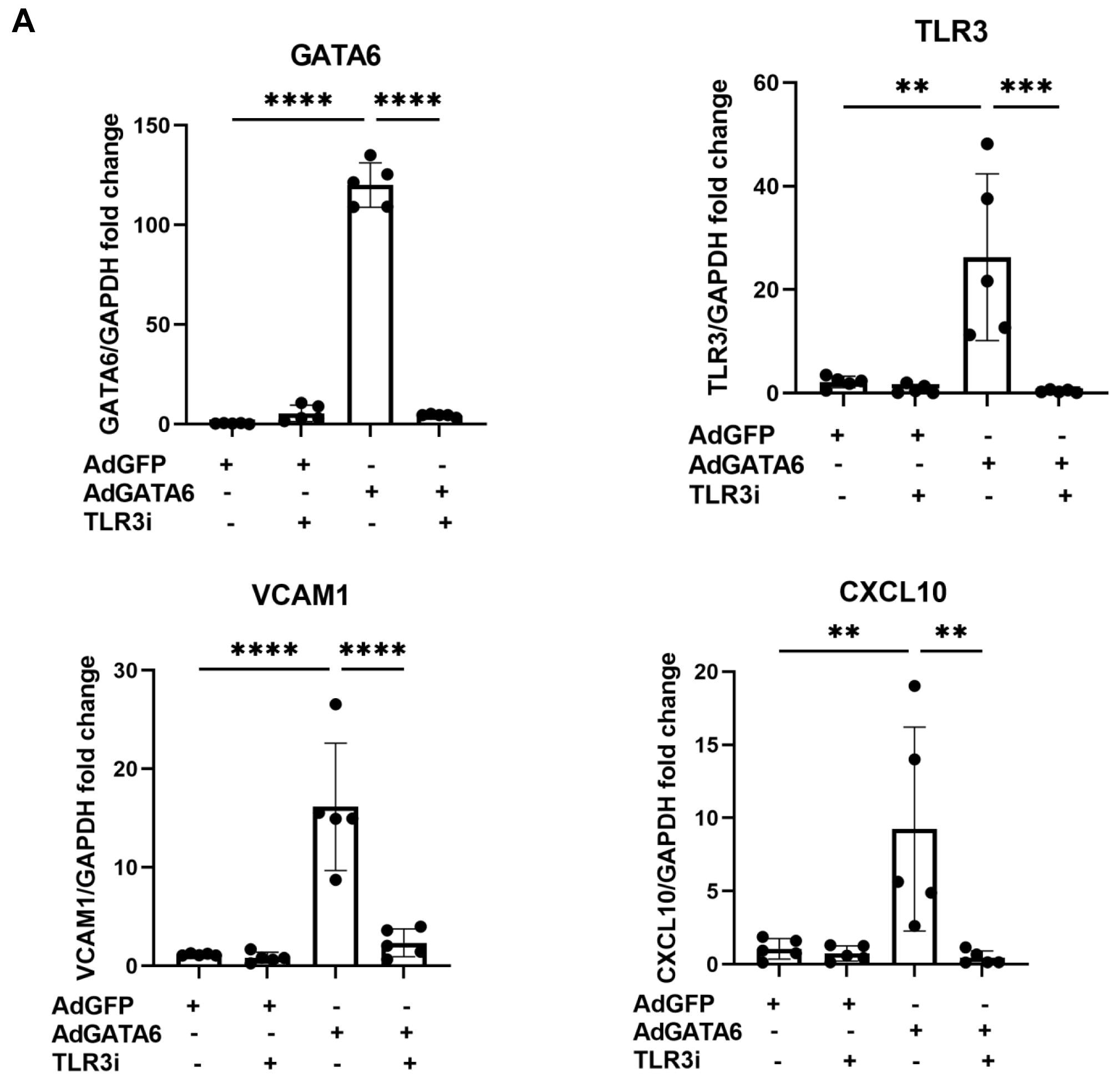
**Inhibition of TLR3 pathway prevents GATA6-dependent induction of interferon genes.** (A) HPAECs were treated with the TLR3 inhibitor (TLR3i, 5 µM) for 6 hours followed by overexpression of GATA6 using AdGATA6 or control AdGFP for 24 hours, and mRNA levels of *GATA6*, *TLR3, VCAM1*, and *CXCL10* were quantified by RT-qPCR. Overexpression of GATA6 markedly upregulated TLR3 and interferon-related targets (VCAM1 and CXCL10), whereas pharmacological inhibition of TLR3 abolished this effect. Data are presented as mean ± SD (n = 5 independent experiments performed in triplicate). p-values were determined using one-way ANOVA with Tukey’s post hoc test. Exact p-values were: *<u>GATA6</u>*: AdGFP(+)/AdGATA6(–)/TLR3i(–) vs AdGFP(–)/AdGATA6(+)/TLR3i(–), p < 0.0001, and AdGFP(–)/AdGATA6(+)/TLR3i(–) vs AdGFP(–)/AdGATA6(+)/TLR3i(+), p < 0.0001; *<u>TLR3</u>*: p = 0.0012, p = 0.0006; *<u>VCAM1</u>*: p < 0.0001, p < 0.0001; *<u>CXCL10</u>*: p = 0.0099, p = 0.0057.

### GATA6 overexpression induces interferon-response genes independent of TLR3 signaling

To further confirm that the effects of GATA6 on interferon response gene expression are mediated through the TLR3 pathway, we performed combined GATA6 overexpression and TLR3 knockdown experiments in HPAECs. Cells were infected with AdGATA6 or control AdGFP, with or without TLR3 siRNA transfection, and the expression of *VCAM1* and *CXCL10* was analyzed by qPCR.

As shown in Figure 5, GATA6 overexpression markedly increased the mRNA levels of TLR3, VCAM1, and CXCL10 compared to control AdGFP-transduced cells. Unexpectedly, siTLR3 did not blunt the AdGATA6-driven response; siTLR3 modestly augmented the induction of *VCAM1* and *CXCL10* (Fig. 5A), despite effectively lowering TLR3 mRNA. Thus, AdGATA6 can elevate these interferon-responsive genes independently of TLR3, and/or TLR3 signaling may impose negative feedback that limits their magnitude under basal conditions. This contrasts with the TLR3/dsRNA complex inhibitor, which reduced AdGATA6-induced transcripts; the difference likely reflects mechanism-specific effects (acute blockade of endosomal dsRNA signaling by the inhibitor versus receptor depletion that may trigger compensatory RIG-I/MDA5 or NF-κB pathways) and suggests that GATA6 amplifies interferon/inflammatory genes through both TLR3-dependent and TLR3-independent routes.

**Figure 5:**
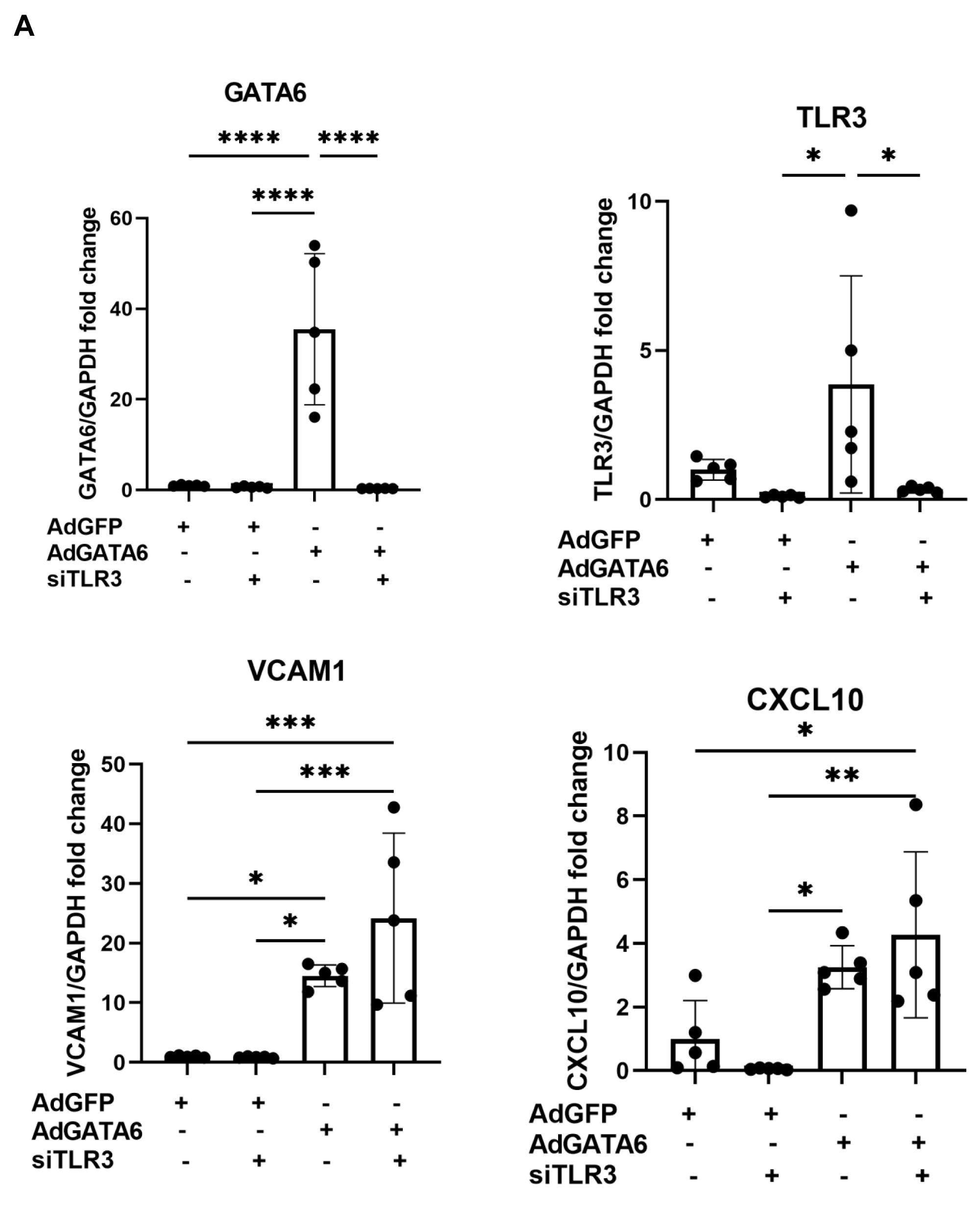
**GATA6 overexpression induces interferon-related genes independent of TLR3 signaling** (A) HPAECs were transfected with siTLR3 or control siRNA for 24 hours, followed by infection with AdGATA6 or control AdGFP for an additional 24 hours, and mRNA levels of *GATA6, TLR3, VCAM1*, and *CXCL10* were quantified by RT-qPCR. Data are presented as mean ± SD (n = 5 independent experiments performed in triplicate). p-values were determined using one-way ANOVA with Tukey’s post hoc test. Exact p-values were as follows: For *<u>GATA6</u>*, AdGFP(+)/AdGATA6(–)/siTLR3(–) vs AdGFP(–)/AdGATA6(+)/siTLR3(–) (p <0.0001), AdGFP(+)/AdGATA6(-)/siTLR3(+) vs AdGFP(–)/AdGATA6(+)/siTLR3(–) (p<0.0001), and AdGFP(–)/AdGATA6(+)/siTLR3(–) vs AdGFP(–)/AdGATA6(+)/siTLR3(+)(p<0.0001). For *<u>TLR3</u>*, AdGFP(+)/AdGATA6(-)/siTLR3(+) vs AdGFP(–)/AdGATA6(+)/siTLR3(–) (p=0.0236), and AdGFP(–)/AdGATA6(+)/siTLR3(–) vs AdGFP(–)/AdGATA6(+)/siTLR3(+) (p=0.0351). For *<u>VCAM1</u>*, AdGFP(+)/AdGATA6(–)/siTLR3(–) vs AdGFP(–)/AdGATA6(+)/siTLR3(–)(p=0.0397),AdGFP(+)/AdGATA6(–)/siTLR3(–)vsAdGFP(–)/AdGATA6(+)/siTLR3(+) (p = 0.0006), AdGFP(+)/AdGATA6(-)/siTLR3(+) vs AdGFP(–)/AdGATA6(+)/siTLR3(–) (p=0.0374), and AdGFP(+)/AdGATA6(-)/siTLR3(+) vs AdGFP(–)/AdGATA6(+)/siTLR3(+) (p=0.0005). For *<u>CXCL10</u>*, AdGFP(+)/AdGATA6(–)/siTLR3(–) vs AdGFP(–)/AdGATA6(+)/siTLR3(+) (p=0.0139), AdGFP(+)/AdGATA6(–)/siTLR3(+) vs AdGFP(–)/AdGATA6(+)/siTLR3(-) (p=0.0165), and AdGFP(+)/AdGATA6(-)/siTLR3(+) vs AdGFP(–)/AdGATA6(+)/siTLR3(+) (p=0.0018).

### Conditioned medium from GATA6-deficient HPAECs upregulates interferon target genes in HPASMCs

We next aimed to investigate the potential paracrine effects of endothelial GATA6 deficiency on neighboring cells within the vascular environment. Given the established interactions between endothelial cells and smooth muscle cells in the vasculature, we hypothesized that the loss of GATA6 in endothelial cells might alter the secretion of soluble factors, thereby influencing immune responses in adjacent smooth muscle cells.

To test this hypothesis, HPAECs were transfected with an siRNA targeting GATA6 or a control scrambled siRNA for 24 hours. The medium was then replaced with serum-free media to generate conditioned medium (CM), which was collected 24 hours later. This CM was used to treat healthy HPASMCs for 24 hours. After treatment, RNA sequencing was performed on the HPASMCs to evaluate changes in gene expression induced by the CM. Gene Set Enrichment Analysis (GSEA) indicated significant (FDR *q* < 0.05) coordinate upregulation of genes involved in interferon signaling (Fig 6A). Selected upregulated genes were further validated using qPCR (Fig 6B). Our results demonstrated that CM from GATA6-deficient HPAECs significantly upregulated *TLR3* as well as interferon target genes including *IFIT1*, *OAS3*, *OAS1*, *MX1* and *IFI44L* in HPASMCs. These findings suggest that GATA6 plays an important role not only in the intrinsic regulation of endothelial immune responses but also in the endothelial-smooth muscle cell crosstalk in the pulmonary vasculature.

**Figure 6:**
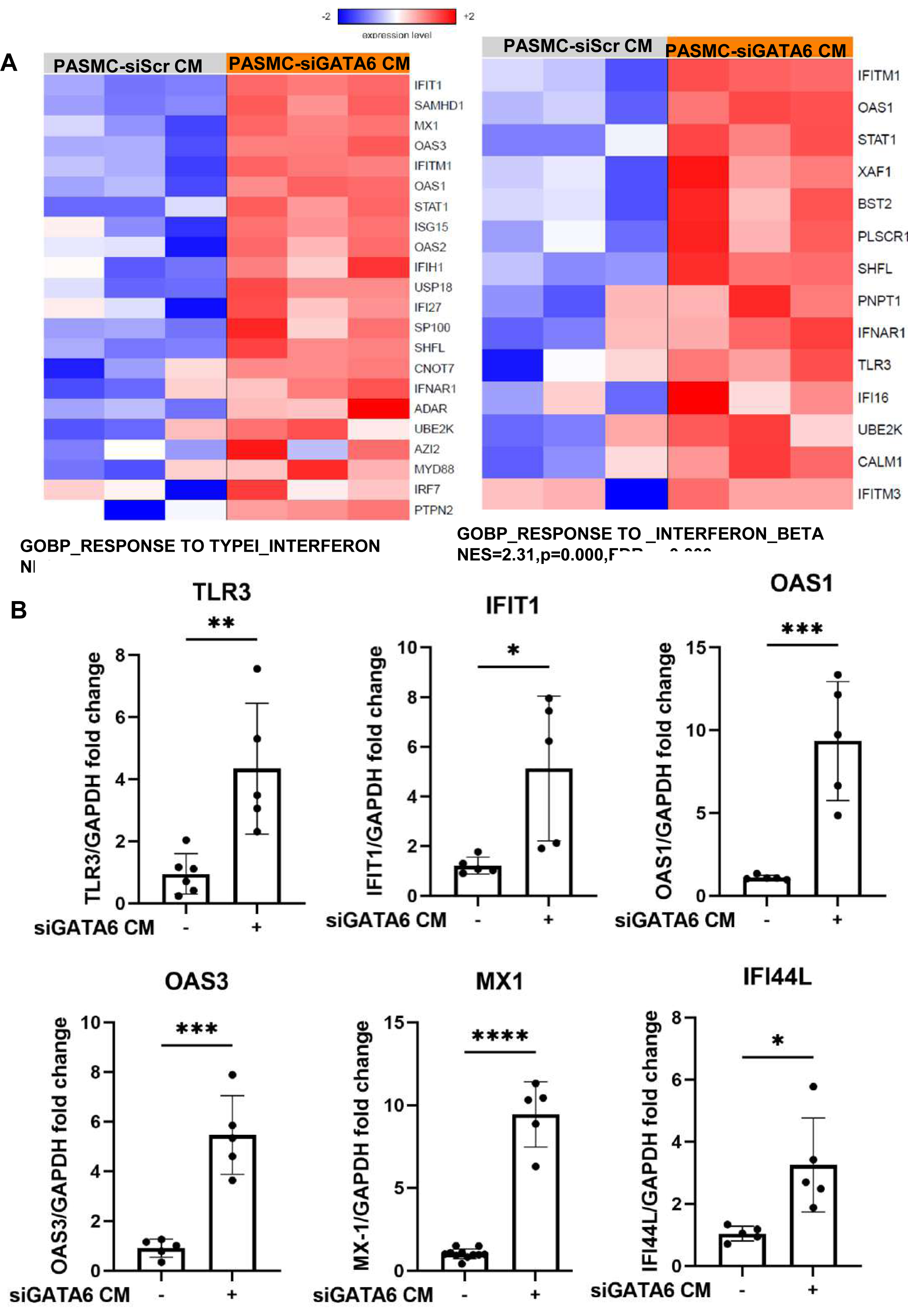
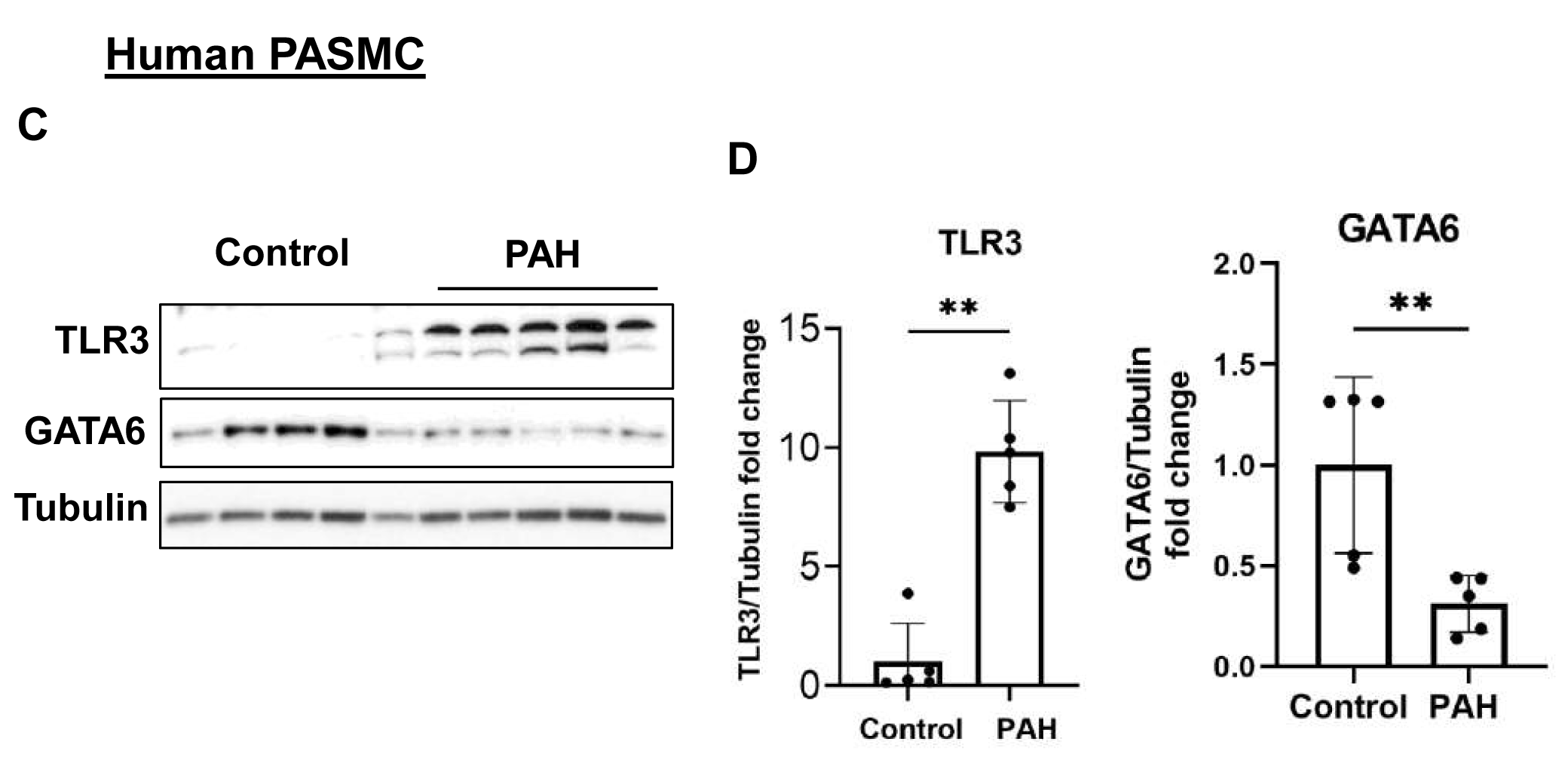
**Conditioned medium from GATA6 deficient HPAECs leads to upregulation of interferon target genes in HPASMCs.** (A) RNA sequencing of HPASMCs treated with conditioned medium (CM) from siScr- or siGATA6-transfected HPAECs revealed enrichment of gene sets associated with type I interferon response (GOBP_RESPONSE_TO_TYPE_I_INTERFERON; NES = 2.33, *p* = 0.000, FDR = 0.000) and interferon-beta signaling (GOBP_RESPONSE_TO_INTERFERON_BETA; NES = 2.31, *p* = 0.000, FDR = 0.000). The heatmaps display relative expression levels of representative interferon-stimulated genes (ISGs). (B) Validation of selected ISGs by RT-qPCR confirmed that CM from GATA6-deficient HPAECs significantly upregulated *TLR3, IFIT1, OAS1, OAS3, MX1*, and *IFI44L* in HPASMCs. Data are means ± SD (n=5, average of five independent experiments done in triplicates), statistical analysis was performed using an unpaired two-tailed Student’s t-test. Exact p-values: *<u>TLR3</u>* (p =0.0044), *<u>IFIT1</u>* (p = 0.0177), *<u>OAS3</u>* (p = 0.0002), *<u>OAS1</u>* (p = 0.0009), *<u>MX1</u>*(p <0.0001) and *<u>IFI44L</u>* (p = 0.0121). (C) PASMCs from non-diseased (control) subjects and patients with PAH were subjected to immunoblot analysis to detect indicated proteins. Data are means±SD; n=5 subjects/group; statistical analysis was performed using Mann Whitney U. Exact p-values: <u>TLR3</u>(p=0.0040), <u>GATA6(</u>p=0.0040)

In line with these findings, our prior study showed that GATA6 expression is reduced in both pulmonary arterial endothelial and smooth muscle cells isolated from patients with PAH.^19^ To further examine the status of TLR3 signaling in these cells, we compared PASMCs from PAH patients and non-diseased controls To answer this question experimentally, we performed comparative analysis of PASMCs from patients with PAH and non-diseased subjects. In agreement with our previously published study human PAH PASMCs had significantly reduced GATA6 protein levels compared to non-diseased controls. Interestingly, in contrast to HPAECs, GATA6 deficiency in PAH PASMCs was accompanied by significant increase in TLR3 protein levels. This new data is in good agreement with our findings showing that media conditioned by GATA6-deficient HPAECs significantly increases TLR3 mRNA in non-diseased HPASMCs (Figure 6C, D, Supplementary Figure 4 https://doi.org/10.6084/m9.figshare.30680609) and suggests that the effects of GATA6 on TLR3 are cell-specific and differ between HPAECs and HPASMCs. To assess potential paracrine effects of GATA6-deficient PASMCs on endothelial interferon signaling, HPAECs were treated with conditioned media from siGATA6- or siScr-transfected PASMCs; however, no significant changes were observed in GATA6, TLR3, or interferon-response gene expression (Supplementary Fig. 6A,B, https://doi.org/10.6084/m9.figshare.30680609).

### RNA components in conditioned medium from GATA6-Deficient HPAECs mediate interferon gene induction in HPASMCs

To determine the nature of the soluble mediators driving the interferon response in HPASMCs, we next treated the CM from GATA6-deficient HPAECs with RNase to selectively degrade extracellular RNA species prior to application on smooth muscle cells. RNase treatment markedly attenuated the CM-induced upregulation of *TLR3, MX1,* and *IFIT1* mRNA in HPASMCs (Fig. 7A-C). The partial but consistent loss of interferon-stimulated gene induction following RNase treatment indicates that RNA molecules within the endothelial secretome contribute to activating interferon signaling in HPASMCs. These findings suggest that GATA6 deficiency promotes the release of RNA species capable of triggering TLR3-mediated antiviral pathways in adjacent vascular smooth muscle cells, providing mechanistic evidence that the endothelial–smooth muscle crosstalk in this context is at least partly mediated by RNA-dependent signaling.

**Figure 7:**
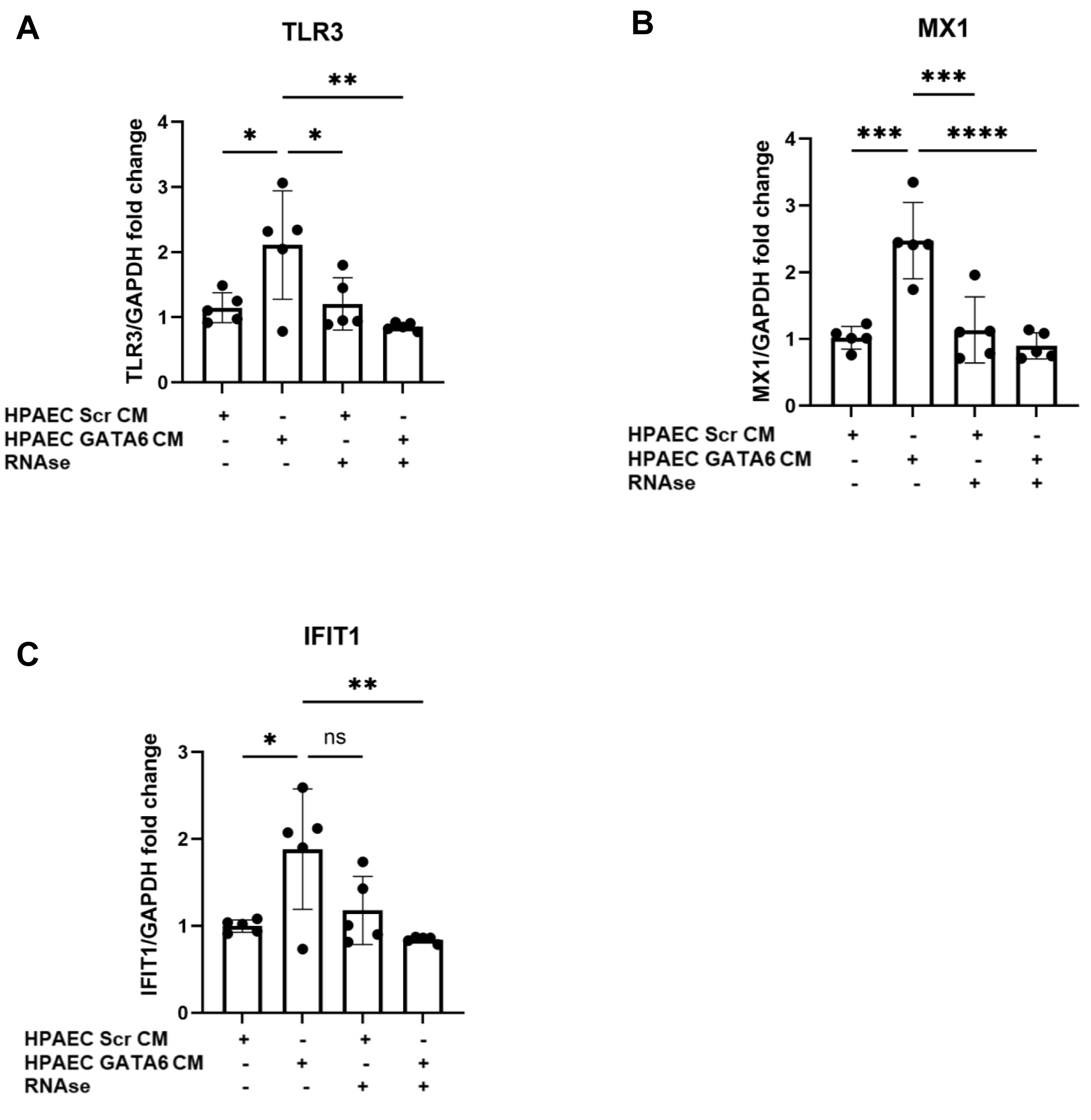
**RNA components in conditioned medium from GATA6-deficient HPAECs mediate interferon gene induction in HPASMCs** (A–C) HPASMCs were treated for 24 hours with conditioned medium (CM) derived from siScr- or siGATA6-transfected HPAECs in the presence or absence of RNase (10µg/mL) to degrade extracellular RNA. Gene expression analysis by RT-qPCR showed that CM from GATA6-deficient HPAECs significantly upregulated *TLR3* (A), *MX1* (B), and *IFIT1* (C) mRNA levels compared with control CM. RNase treatment completely abolished this upregulation. Data are presented as mean ± SD (n = 5 independent experiments performed in triplicate). p-values were calculated using one-way ANOVA with Tukey’s post hoc test. Exact p-values for all pairwise comparisons were as follows: For *<u>TLR3</u>*, HPAEC Scr CM(+) / HPAEC GATA6 CM(–) / RNase(–) vs HPAEC Scr CM(–) / HPAEC GATA6 CM(+) / RNase(–) (p = 0.0262), HPAEC Scr CM(–) / HPAEC GATA6 CM(+) / RNase(–) vs HPAEC Scr CM(+) / HPAEC GATA6 CM(-) / RNase(+) (p = 0.0386), and HPAEC Scr CM(-) / HPAEC GATA6 CM(+) / RNase(–) vs HPAEC Scr CM(–) / HPAEC GATA6 CM(+) / RNase(+) (p = 0.0037). For *<u>MX1</u>*, HPAEC Scr CM(+) / HPAEC GATA6 CM(–) / RNase(–) vs HPAEC Scr CM(–) / HPAEC GATA6 CM(+) / RNase(–) (p = 0.0002), HPAEC Scr CM(–) / HPAEC GATA6 CM(+) / RNase(–) vs HPAEC Scr CM(+) / HPAEC GATA6 CM(-) / RNase(+) (p = 0.0004) , HPAEC Scr CM(-) / HPAEC GATA6 CM(+) / RNase(–) vs HPAEC Scr CM(–) / HPAEC GATA6 CM(+) / RNase(+) (p <0.0001). For *<u>IFIT1</u>*, HPAEC Scr CM(+) / HPAEC GATA6 CM(–) / RNase(–) vs HPAEC Scr CM(–) / HPAEC GATA6 CM(+) / RNase(–) (p = 0.0140), HPAEC Scr CM(–) / HPAEC GATA6 CM(+) / RNase(–) vs HPAEC Scr CM(+) / HPAEC GATA6 CM(-) / RNase(+) (p = ns) , HPAEC Scr CM(-) / HPAEC GATA6 CM(+) / RNase(–) vs HPAEC Scr CM(–) / HPAEC GATA6 CM(+) / RNase(+) (p =0.0040)

## Discussion

Endothelial cells are the first line of defense of the blood vessel and play a crucial role in regulating immune response. Dysregulation in EC function can occur due to cytokine production via pattern recognition receptors (PRRs), triggering innate immune signaling and contributing to a proinflammatory signaling cascade. Several cytokines such as IL-13 and the chemokine CX3CL1, have been implicated in promoting PASMC proliferation in small muscularized PAs, contributing to vascular remodeling in PAH ^27,28^ .

PAH has been increasingly linked to interferon (IFN) therapy. The interferons are signaling proteins central to the body’s immune response against pathogens, including viruses, bacteria, or tumor cells. Type I interferons (IFN-α, IFN-β) have been implicated in PAH development. For example, IFN-α treatment in hepatitis C patients increases Endothelin-1 (ET-1), a well-known pro-PAH factor.^29^. Similarly, high interferon levels in HIV patients are associated with an increased PAH risk.^30^ Previous studies have shown that patients with systemic sclerosis-associated PAH (SSc-PAH) exhibit higher levels of IFN-α and IFNγ in serum compared to SSc patients without PAH. Additionally, ET-1 and pro-inflammatory cytokine CXCL10, along with interferon-inducible genes, are significantly upregulated in the serum of SSc-PAH patients.^31^

Our previous research revealed the role of GATA6 in regulating oxidative stress and BMP10 signaling in’ ECs and SMCs of both SSc-PAH and idiopathic PAH (IPAH) patients.^19^ This study further demonstrates that GATA6 deficiency, confirmed through both siRNA-mediated depletion and the use of pyrrothiogatain (a pan-GATA inhibitor), results in significant downregulation of interferon response genes and TLR3 expression in HPAECs. While increase in pro-inflammatory signaling is commonly associated with PAH progression, Farkas et al have reported a loss of TLR3 expression in the lungs and PAECs of PAH patients. Additionally, they demonstrated that TLR3 deficient PAECs mediate RNA signaling through cytoplasmic RNA sensors RIG-1 and MDA5, inducing apoptosis.^5^ Our data further show that along with TLR3 expression levels, RIG-1 and MDA5 mRNA levels are also downregulated in GATA6 deficient cells, suggesting that GATA6 deficiency disrupts RNA sensing by reducing TLR3, MDA5 and RIG-1 expression, thus impacting the associated immune response.

Previous studies have demonstrated that TLR3 induction via PolyIC exerts a protective role in the arteries by reducing neointima formation.^12^ Additionally, PolyIC treatment reduced PH, improved right ventricular function, and attenuated proliferation and apoptosis in pulmonary artery in a chronic hypoxia/Sugen rat model^5^. Here, we show that the GATA inhibitor attenuated the PolyIC-induced response of TLR3 and associated interferon genes, VCAM1 and CXCL10. Activation of TLR3 by the viral RNA mimic Poly(I:C) increased GATA6 protein levels, suggesting a feedback loop wherein TLR3 signaling stabilizes or enhances GATA6 expression to reinforce interferon responses. Conversely, inhibition of GATA6 attenuated Poly(I:C)-induced upregulation of TLR3, CXCL10, and VCAM1, indicating that GATA6 is required for full activation of this antiviral program.Notably, PolyIC stimulation of HPAECs has been shown to increase BMPR2 expression by promoting IRF3 binding to the BMPR2 promoter.^32^ Our earlier work has demonstrated that GATA6 also contributes to BMPR2 expression in HPAECs^19^, suggesting a shared signaling pathway between GATA6 and TLR3/IRF3 in regulating expression of BMPR2.This regulatory relationship highlights the importance of GATA6 in modulating the TLR3 receptor expression, its downstream signaling pathway, and the associated interferon response in HPAECs, which may have implications for pulmonary vascular diseases and responses to viral infections. Overexpression studies further revealed that GATA6 amplifies interferon-related gene expression through both TLR3-dependent and -independent mechanisms. While adenoviral GATA6 increased TLR3, VCAM1, and CXCL10 expression, these effects were abolished by pharmacologic inhibition of TLR3 signaling. However, siRNA-mediated depletion of TLR3 unexpectedly enhanced the induction of VCAM1 and CXCL10 by GATA6 overexpression, suggesting that TLR3 may exert a negative feedback role under certain basal conditions. Together, these findings imply that GATA6 acts both upstream of TLR3 to promote receptor expression and downstream or parallel to TLR3 to sustain interferon-responsive transcription, depending on the cellular context and feedback state of the pathway.

Beyond its cell-autonomous effects, GATA6 deficiency profoundly altered the endothelial secretome, as evidenced by RNA sequencing of HPASMCs exposed to conditioned media from siGATA6-transfected HPAECs. Smooth muscle cells treated with this conditioned medium exhibited marked upregulation of TLR3, IFIT1, MX1, OAS1, and other interferon response genes, indicating that endothelial GATA6 loss triggers paracrine signaling that activates antiviral and inflammatory pathways in neighboring cells. RNase treatment of conditioned media attenuated this effect, implicating extracellular RNA species as major mediators of endothelial–smooth muscle communication. Furthermore, inhibition of GATA6 blunted Poly(I:C)-induced IFNα/β production, indicating that GATA6 activity is required for optimal interferon pathway activation. These findings provide mechanistic evidence that GATA6 deficiency promotes the release of RNA and cytokine mediators capable of engaging TLR3 signaling in adjacent smooth muscle cells.

These findings provide mechanistic evidence that GATA6 deficiency enhances endothelial extracellular RNA mediated signaling and interferon pathway dysregulation, collectively driving paracrine activation of adjacent smooth muscle cells through TLR3-related mechanisms. Interestingly, while GATA6 deficiency downregulates TLR3/interferon signaling in HPAECs, conditioned medium from GATA6-deficient cells induced interferon response genes in PASMCs. In PASMCs from PAH patients, reduced GATA6 coincides with increased TLR3 protein expression. The role of interferon pathway in PASMCs in the context of PAH remains complex, though evidence suggests that elevated interferon signaling may contribute to vascular remodeling, inflammation, and metabolic dysregulation. ^31^ Relevant to our findings, single-cell RNA sequencing of PAH PASMCs identified four transcriptionally distinct clusters that showed differential distribution in healthy and PAH lungs. The synthetic cluster of PASMCs, characterized by Type I interferon response genes, as well as genes related to ECM deposition and cell migration, was enriched in patients’ lungs.^33^

The functional proliferative consequences of endothelial GATA6 deficiency have already been demonstrated by our group^19^ and the present work advances this framework by identifying interferon pathway activation as a complementary mechanism of endothelial-to-SMC communication in pulmonary vascular remodeling. In summary, these data support a model in which GATA6 imposes cell-type–specific transcriptional control across the vascular wall. Loss of endothelial GATA6 not only weakens intrinsic antiviral defenses but also promotes a proinflammatory paracrine milieu that activates interferon pathways in PASMCs together providing a coherent mechanistic link between endothelial dysfunction and smooth muscle remodeling in PAH. Future studies using defined co-culture or organ-on-chip systems will be important to map GATA6-dependent vesicular/RNA secretory pathways with higher resolution, and to determine whether targeted restoration of endothelial GATA6 can re-establish vascular immune homeostasis.

## Supplemental Data

Bulk RNA-sequencing data have been deposited in GEO under accession number GSE293379. All Supplemental Data, including Supplementary Figures and Supplementary Table 1 containing the full GSEA results for all comparisons, are available at: https://doi.org/10.6084/m9.figshare.30680609.

## Data Availability

The full set of gene expression measurements from the microarray experiment mentioned in this study have been previously published (Supplemental Table 1) [PMID 37087509]. The raw and normalized RNA-seq data have been deposited in the Gene Expression Omnibus (GEO), Series GSE293379.

## Acknowledgments

The authors thank Dr. Francesca Seta and Dr. Andreea Bujor at Boston University Chobanian & Avedisian School of Medicine for their valuable guidance and support throughout this study. Their expertise and insightful feedback greatly contributed to the quality and rigor of this work. The graphical abstract was made by using Biorender.com

Grants

The work is supported by *NIH/National Heart, Lung, and Blood Institute* 5R01HL150638-04(M.T), R01 HL172488(E.A.G).

## Disclosures

The authors report no conflicts of interest in this work

## Author Contributions

Conception and design: V.D, M.T; Experimental work: V.D,I.Za,D.G,T.D,I.Zh ; Analysis of data and preparation of figures: V.D, I.Za,T.D,I.Zh ; Interpretation: V.D, M.T,D.G, E.A.G. ; Drafting, editing and revision of the manuscript: V.D, M.T, E.A.G ; Approved final version : V.D,M.T, E.A.G

---

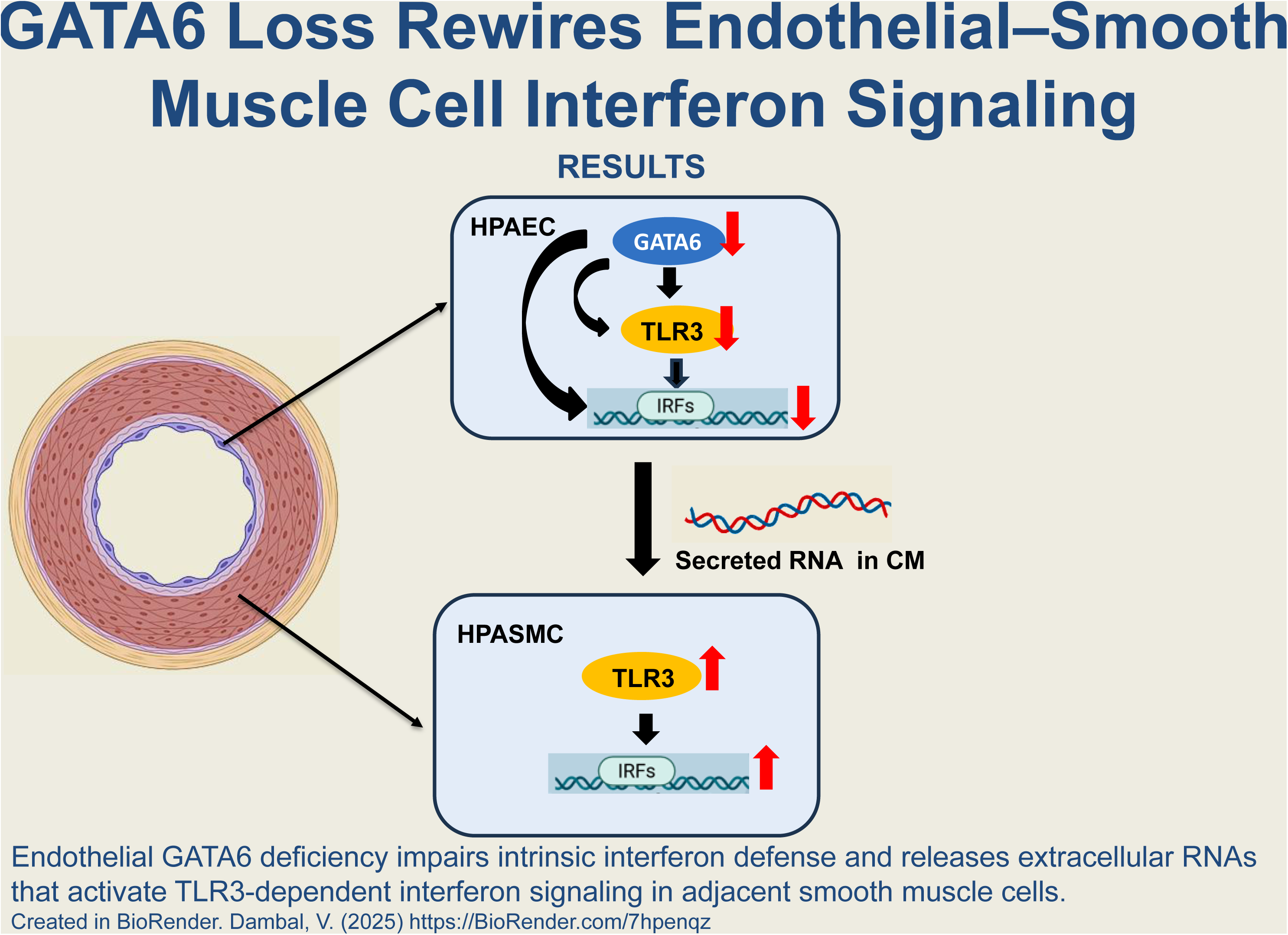

## Notes

https://doi.org/10.6084/m9.figshare.30680609

### Competing Interest Statement

The authors have declared no competing interest.

## References

1. Pober JS, Sessa WC. Evolving functions of endothelial cells in inflammation. Nat Rev Immunol. 2007;7(10):803–15.

2. Dauphinee SM, Karsan A. Lipopolysaccharide signaling in endothelial cells. Lab Invest. 2006;86(1):9–22.

3. Liao JK. Linking endothelial dysfunction with endothelial cell activation. J Clin Invest. 2013;123(2):540–1.

4. Al-Soudi A, Kaaij MH, Tas SW. Endothelial cells: From innocent bystanders to active participants in immune responses. Autoimmun Rev. 2017;16(9):951–62.

5. Farkas D, Thompson AR, Bhagwani AR, et al. Toll-like receptor 3 is a therapeutic target for pulmonary hypertension. Am J Respir Crit Care Med. 2019;199(2):199–210.

6. Alexopoulou L, Holt AC, Medzhitov R, Flavell RA. Recognition of double-stranded RNA and activation of NF-κB by Toll-like receptor 3. Nature. 2001;413(6857):732–8.

7. Lai Y, Yi G, Chen A, et al. Viral double-strand RNA-binding proteins can enhance innate immune signaling by Toll-like receptor 3. PLoS One. 2011;6(10):e25837.

8. Bernard JJ, Cowing-Zitron C, Nakatsuji T, et al. Ultraviolet radiation damages self noncoding RNA and is detected by TLR3. Nat Med. 2012;18(8):1286–90.

9. Zhang SY, Jouanguy E, Ugolini S, et al. TLR3 deficiency in patients with herpes simplex encephalitis. Science. 2007;317(5844):1522–7.

10. Kawai T, Akira S. TLR signaling. Cell Death Differ. 2006;13(5):816–25.

11. Pryshchep O, Ma-Krupa W, Younge BR, Goronzy JJ, Weyand CM. Vessel-specific Toll-like receptor profiles in human medium and large arteries. Circulation. 2008;118(12):1276–84.

12. Cole JE, Navin TJ, Cross AJ, et al. Unexpected protective role for Toll-like receptor 3 in the arterial wall. Proc Natl Acad Sci U S A. 2011;108(6):2372–7.

13. Molkentin JD. The zinc finger-containing transcription factors GATA-4, −5, and −6: Ubiquitously expressed regulators of tissue-specific gene expression. J Biol Chem. 2000;275(50):38949–52.

14. Umetani M, Mataki C, Minegishi N, et al. Function of GATA transcription factors in induction of endothelial vascular cell adhesion molecule-1 by tumor necrosis factor-α. Arterioscler Thromb Vasc Biol. 2001;21(6):917–22.

15. German Z, Chambliss KL, Pace MC, Arnet UA, Lowenstein CJ, Shaul PW. Molecular basis of cell-specific endothelial nitric-oxide synthase expression in airway epithelium. J Biol Chem. 2000;275(11):8183–9.

16. Dorfman DM, Wilson DB, Bruns GA, Orkin SH. Human transcription factor GATA-2: Evidence for regulation of preproendothelin-1 gene expression in endothelial cells. J Biol Chem. 1992;267(2):1279–85.

17. Losa M, Latorre V, Andrabi M, et al. A tissue-specific, GATA6-driven transcriptional program instructs remodeling of the mature arterial tree. eLife. 2017;6:e31362.

18. Ghatnekar A, Chrobak I, Reese C, et al. Endothelial GATA-6 deficiency promotes pulmonary arterial hypertension. Am J Pathol. 2013;182(6):2391–406.

19. Toyama T, Kudryashova TV, Ichihara A, et al. GATA6 coordinates cross-talk between BMP10 and oxidative stress axis in pulmonary arterial hypertension. Sci Rep. 2023;13(1):6593.

20. Goncharov DA, Kudryashova TV, Ziai H, et al. mTORC2 coordinates pulmonary artery smooth muscle cell metabolism, proliferation, and survival in pulmonary arterial hypertension. Circulation. 2014;129(8):864–74.

21. Dobin A, Davis CA, Schlesinger F, et al. STAR: Ultrafast universal RNA-seq aligner. Bioinformatics. 2013;29(1):15–21.

22. Love MI, Huber W, Anders S. Moderated estimation of fold change and dispersion for RNA-seq data with DESeq2. Genome Biol. 2014;15:550.

23. Subramanian A, Tamayo P, Mootha VK, et al. Gene set enrichment analysis: A knowledge-based approach for interpreting genome-wide expression profiles. Proc Natl Acad Sci U S A. 2005;102(43):15545–50.

24. Ghatnekar A, Trojanowska M. GATA-6 is a novel transcriptional repressor of the human Tenascin-C gene expression in fibroblasts. Biochim Biophys Acta Gene Regul Mech. 2008;1779(3):145–51.

25. Nomura S, Takahashi H, Suzuki J, Kuwahara M, Yamashita M, Sawasaki T. Pyrrothiogatain acts as an inhibitor of GATA family proteins and inhibits Th2 cell differentiation in vitro. Sci Rep. 2019;9(1):17335.

26. Cheng K, Wang X, Yin H. Small-molecule inhibitors of the TLR3/dsRNA complex. J Am Chem Soc. 2011;133(11):3764–7.

27. Hecker M, Zasłona Z, Kwapiszewska G, et al. Dysregulation of the IL-13 receptor system: A novel pathomechanism in pulmonary arterial hypertension. Am J Respir Crit Care Med. 2010;182(6):805–18.

28. Perros F, Dorfmüller P, Souza R, et al. Fractalkine-induced smooth muscle cell proliferation in pulmonary hypertension. Eur Respir J. 2007;29(5):937–43.

29. Dhillon S, Kaker A, Dosanjh A, Japra D, Vanthiel DH. Irreversible pulmonary hypertension associated with the use of interferon-α for chronic hepatitis C. Dig Dis Sci. 2010;55(6):1785–90.

30. Speich R, Jenni R, Opravil M, Pfab M, Russi EW. Primary pulmonary hypertension in HIV infection. Chest. 1991;100(5):1268–71.

31. George PM, Oliver E, Dorfmuller P, et al. Evidence for the involvement of type I interferon in pulmonary arterial hypertension. Circ Res. 2014;114(4):677–88.

32. Bhagwani AR, Ali M, Piper B, et al. A p53-TLR3 axis ameliorates pulmonary hypertension by inducing BMPR2 via IRF3. iScience. 2023;26(2):106155.

33. Crnkovic S, Valzano F, Fließer E, et al. Single-cell transcriptomics reveals skewed cellular communication and phenotypic shift in pulmonary artery remodeling. JCI Insight. 2022;7(20):e161892.

