## Supplementary figures and images for "Endothelial *GATA6* deficiency suppresses intracellular TLR3-interferon signaling in HPAECs and promotes interferon response in HPASMCs"

### Supplemental Figure 1

Supplementary Figure 1

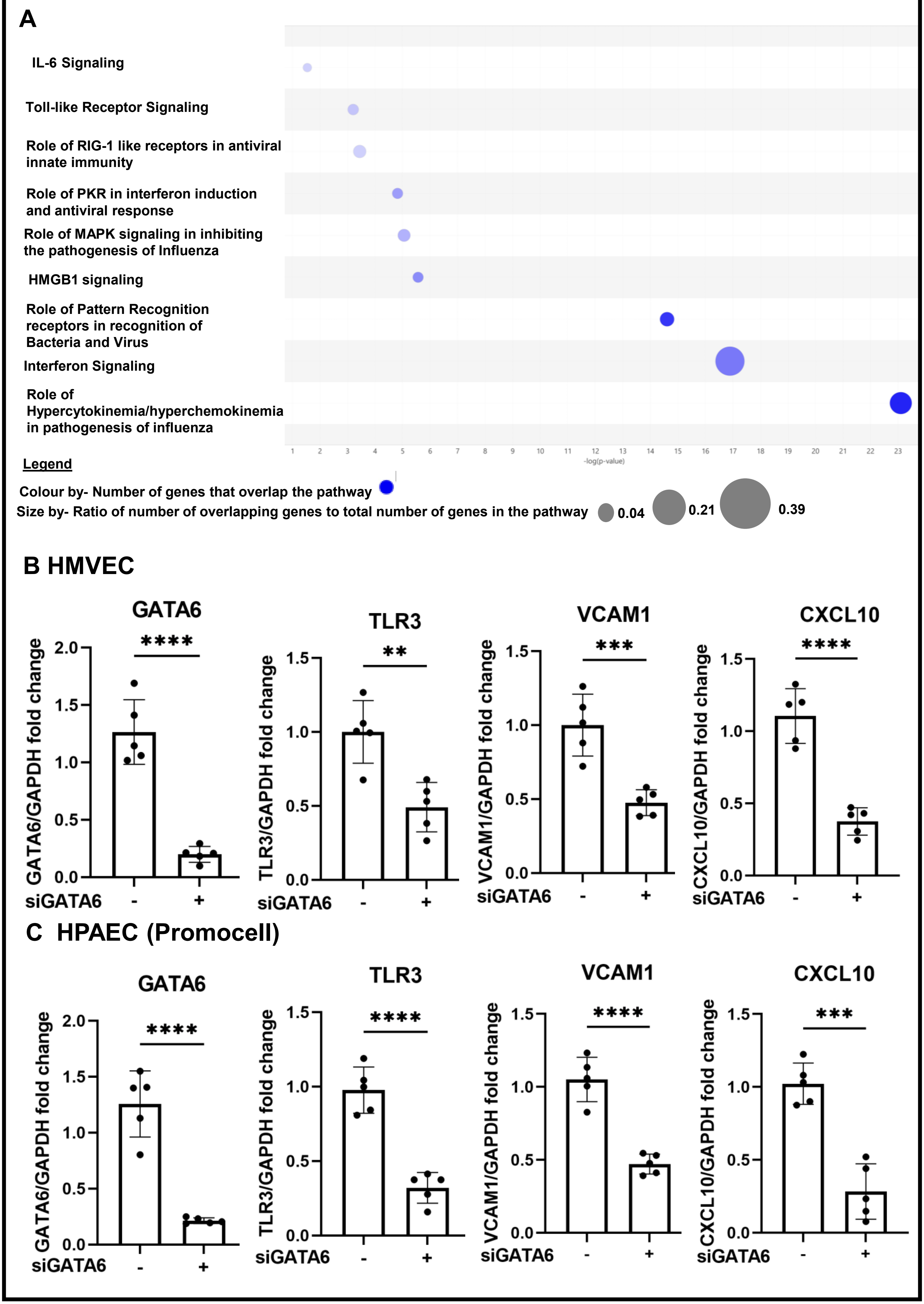

**Supplementary Figure 1**

**D Original Immunoblots for Figure 1**

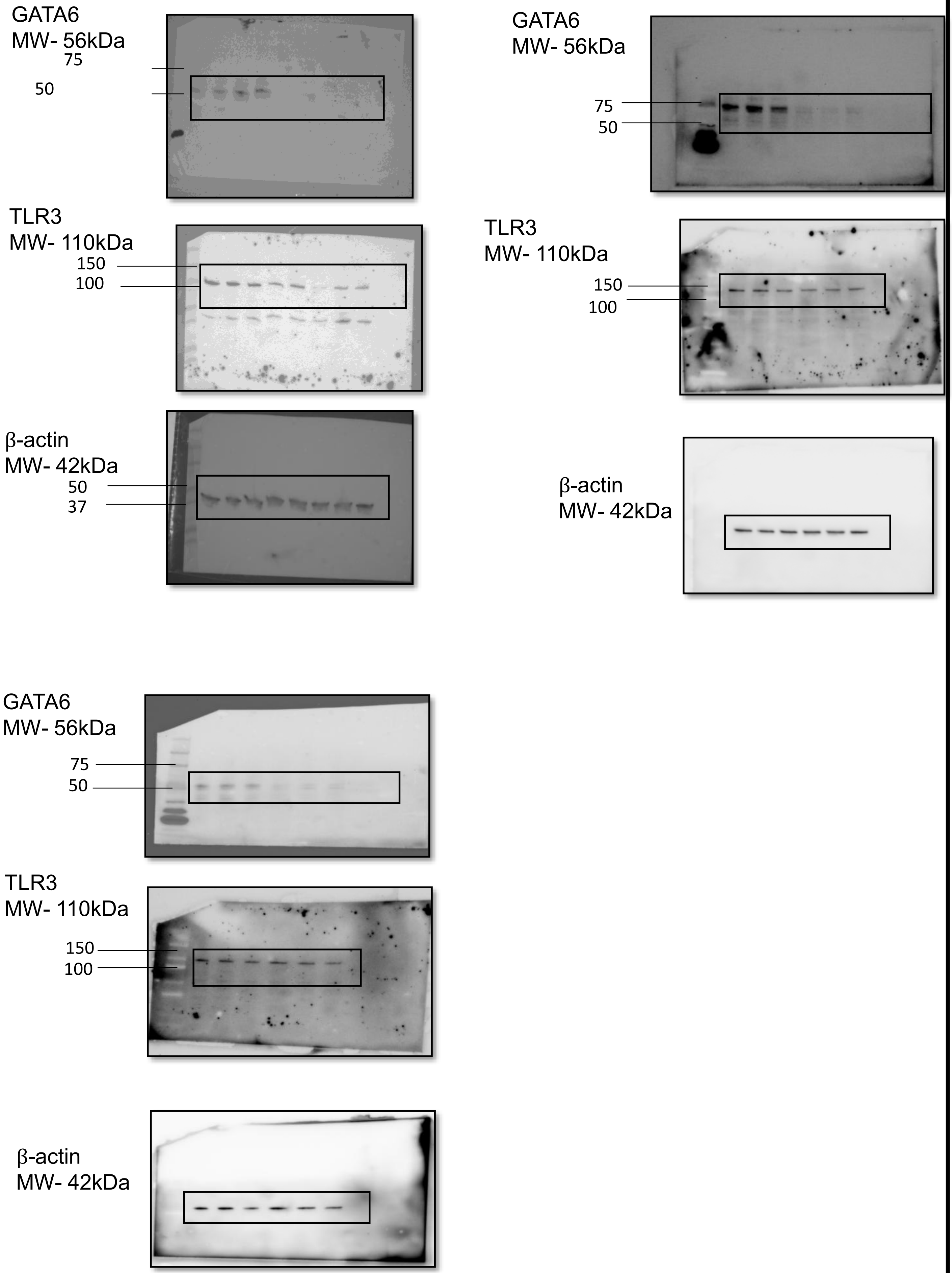

### Supplemental Figure 4

**Supplementary Figure 4**

**A. Original immunoblots for Figure 6**

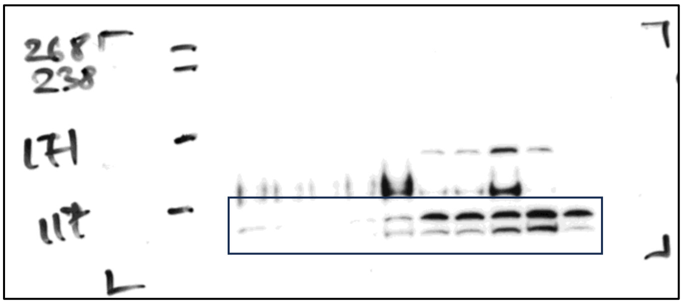


TLR3

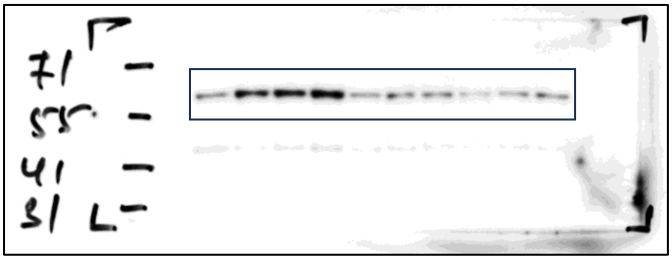


GATA6

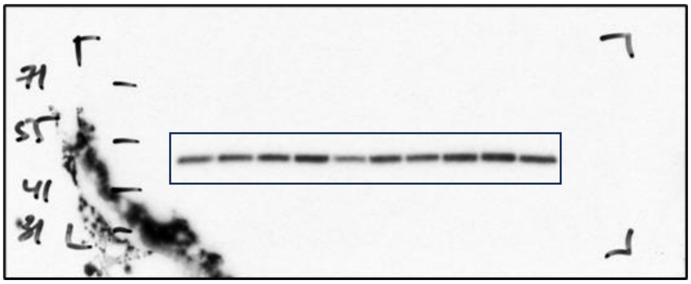


Tubulin

### Supplemental Figure 5

#### Supplementary Figure 5

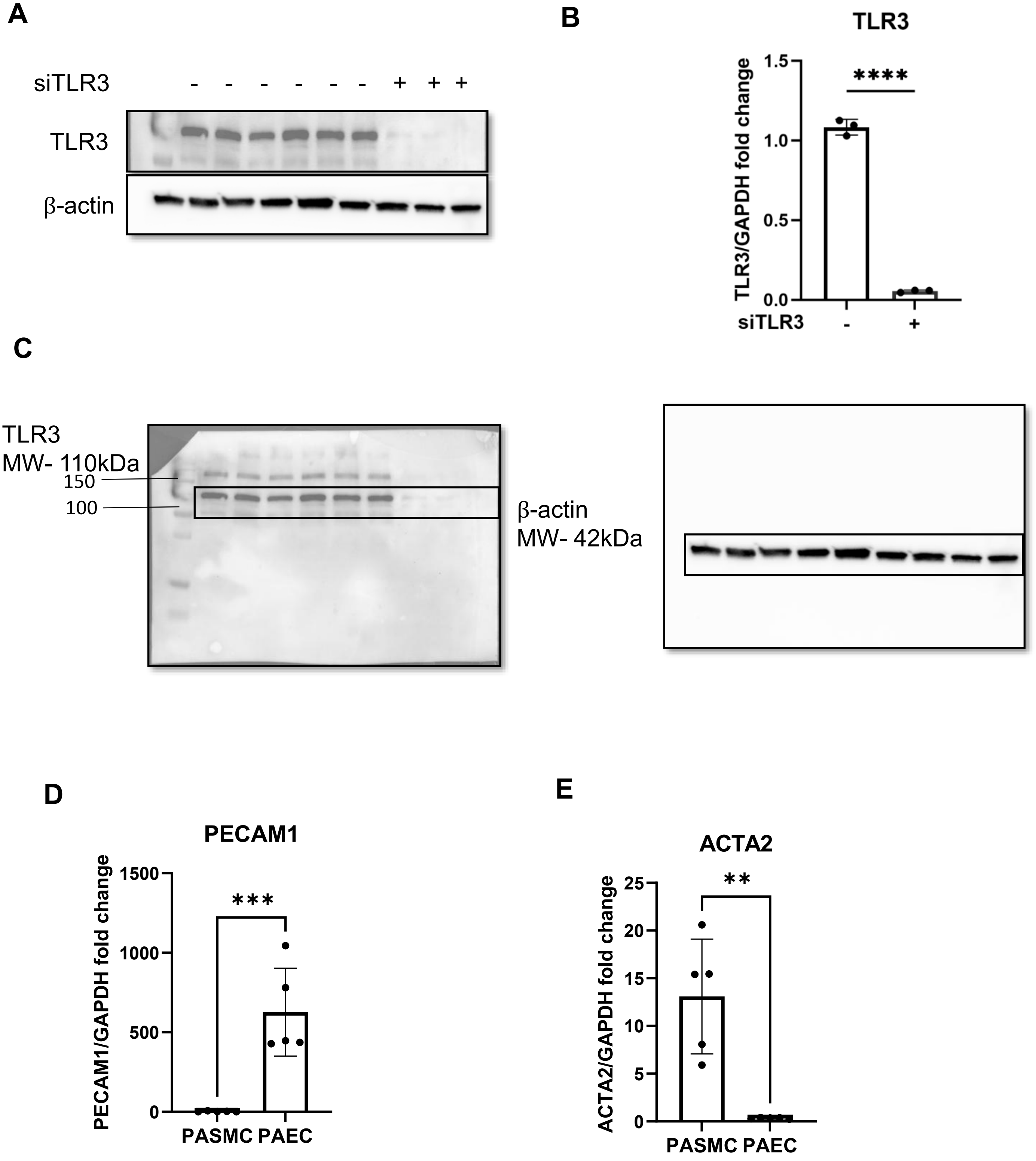

### Supplemental Figure 6

#### Supplementary Figure 6

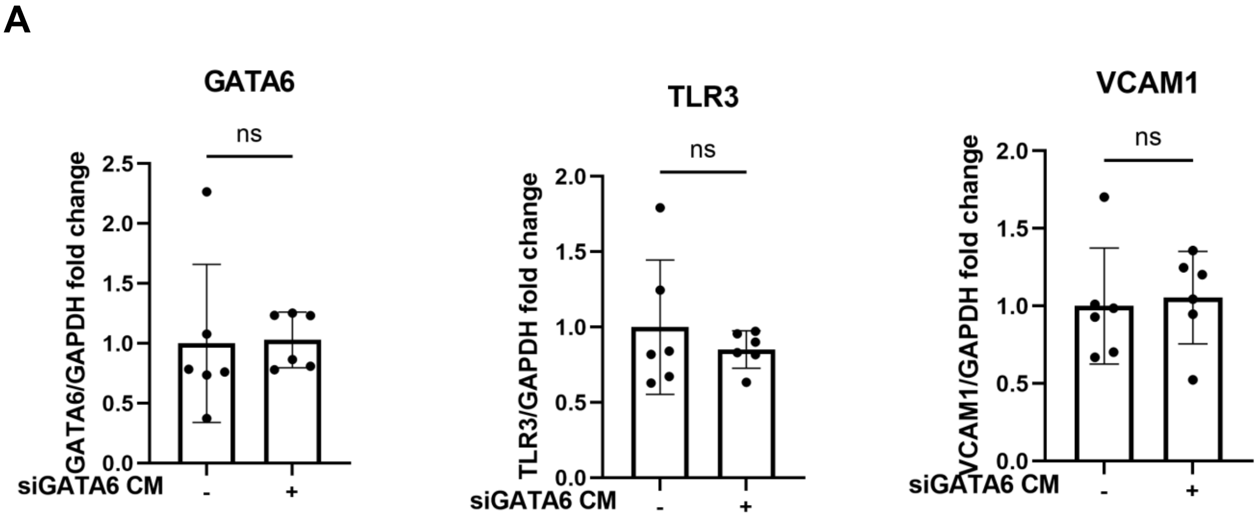


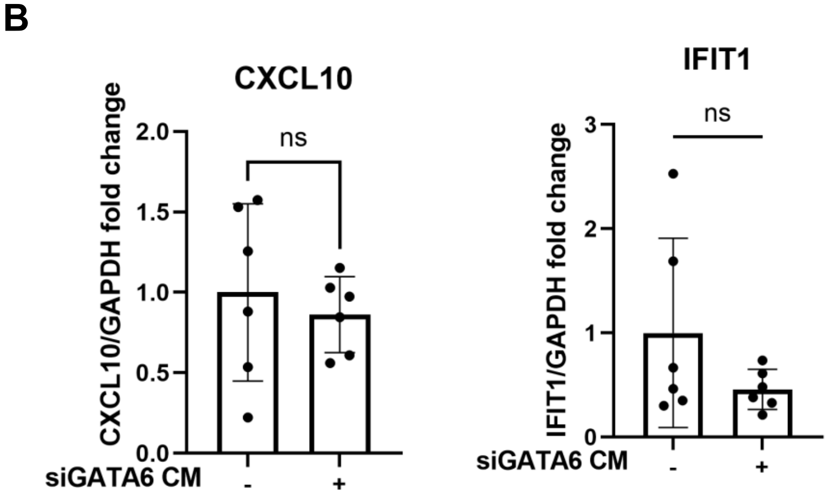
