## Supplemental Figure 2 for "Endothelial *GATA6* deficiency suppresses intracellular TLR3-interferon signaling in HPAECs and promotes interferon response in HPASMCs"

### Supplementary Figure 2

#### A HMVEC

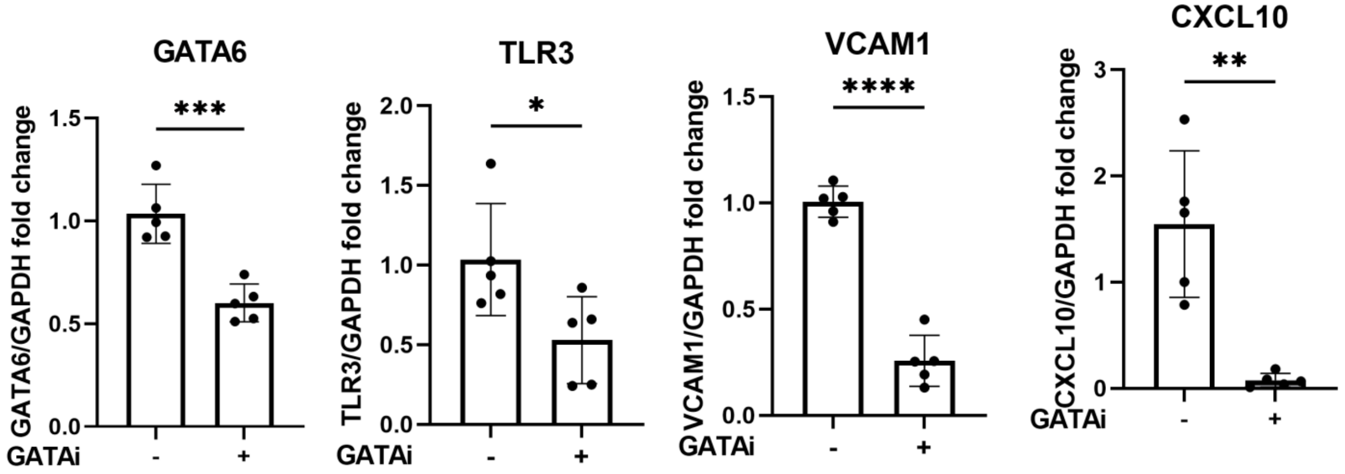


#### B HPAEC (Promocell)

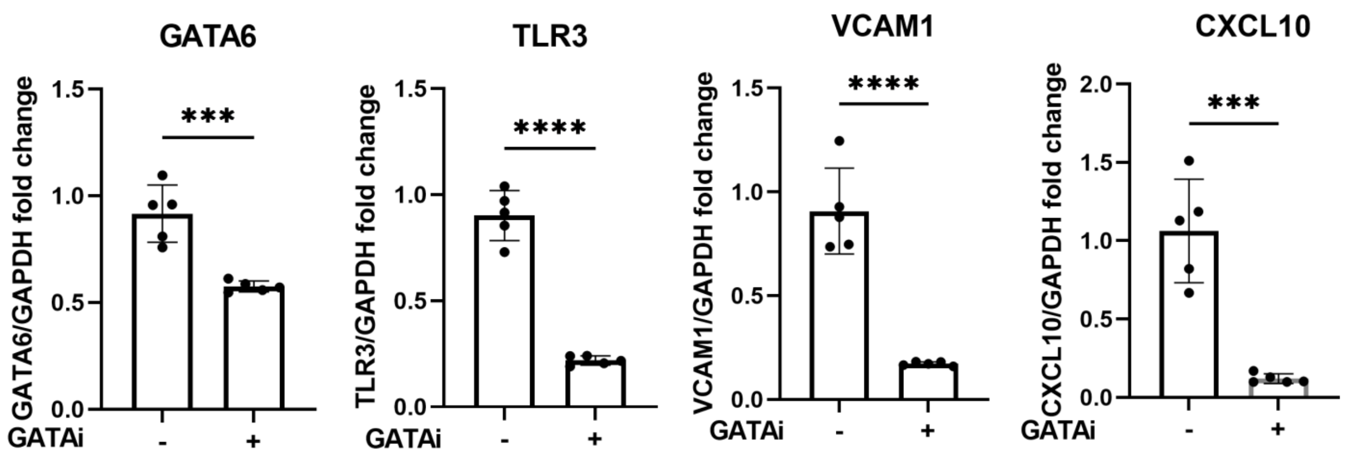


## C

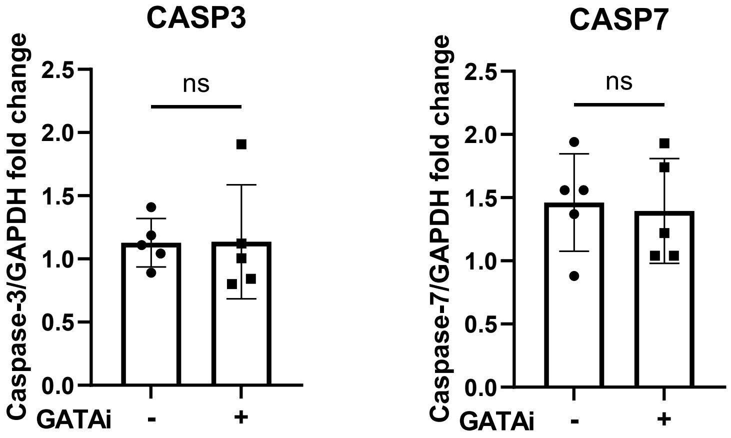

### Supplementary Figure 2

#### D Original Immunoblots for Figure 2

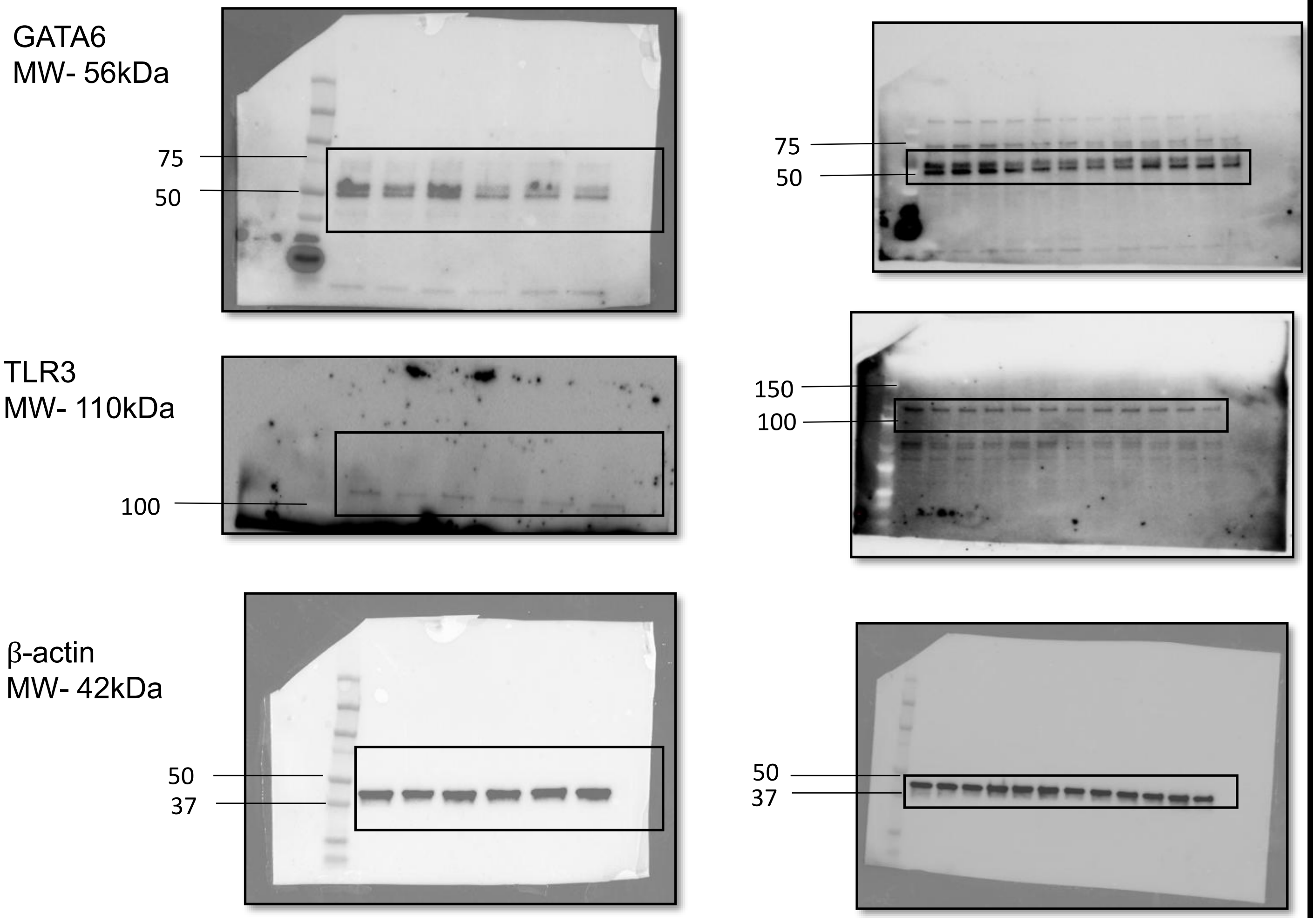
