## Supplemental Figure 3 for "Endothelial *GATA6* deficiency suppresses intracellular TLR3-interferon signaling in HPAECs and promotes interferon response in HPASMCs"

### Supplementary Figure 3

#### A HMVEC

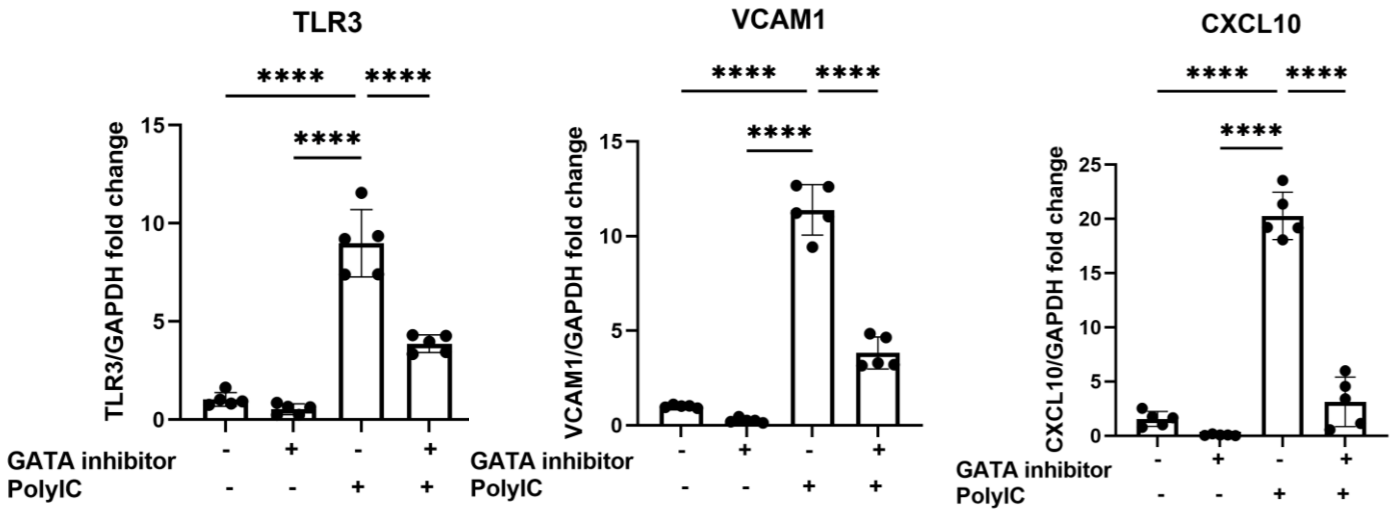


#### B. HPAEC (Promocell)- no TLR3 induction

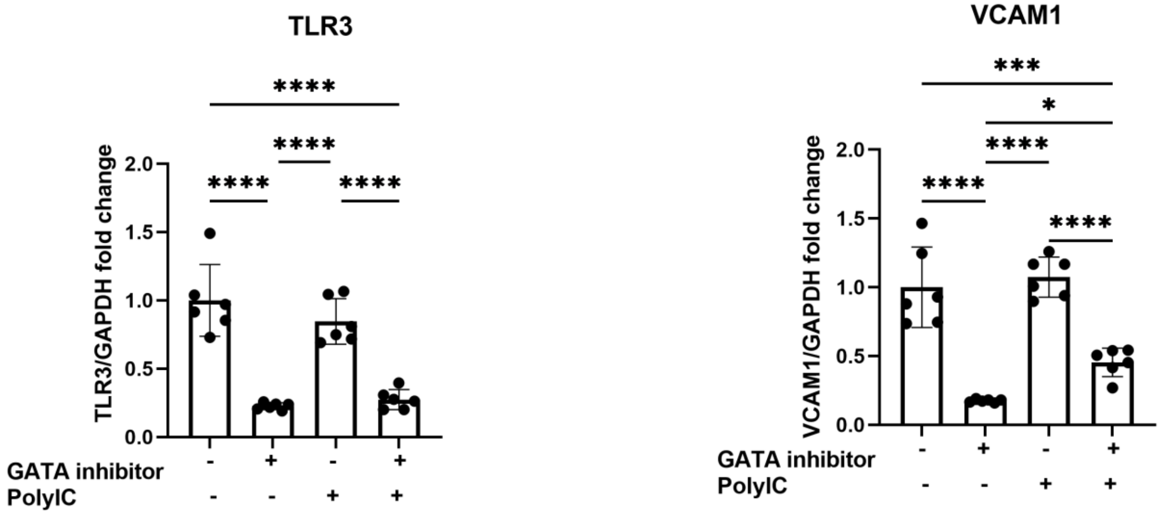

### Supplementary Figure 3

### C Original Immunoblots for Figure 3

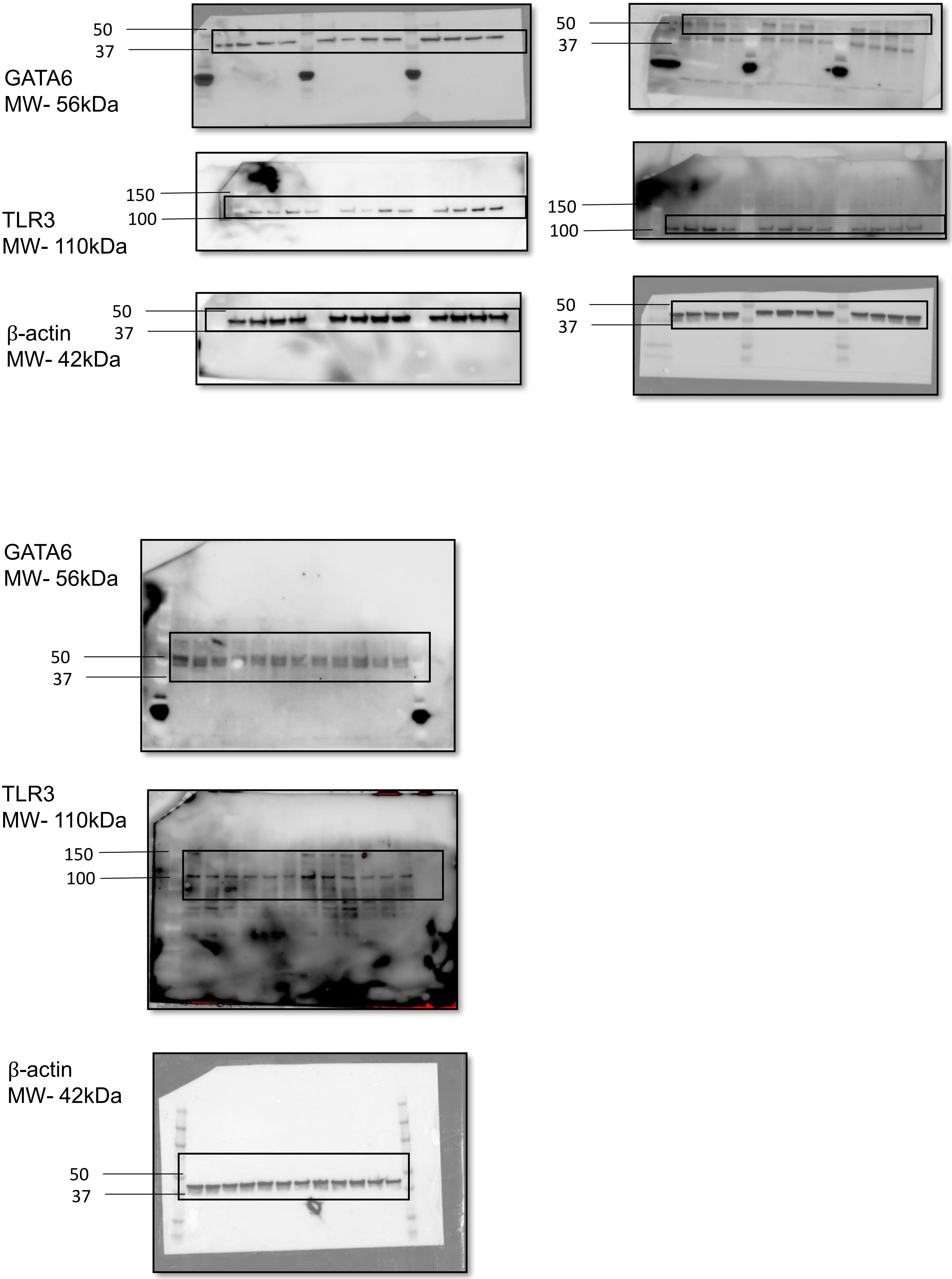
