## Supplemental Figure legend for "Endothelial *GATA6* deficiency suppresses intracellular TLR3-interferon signaling in HPAECs and promotes interferon response in HPASMCs"

**Supplementary Figure legend**

**Supplementary Figure 1**: **GATA6 deficiency in HPAECs leads to downregulation of interferon target genes and TLR3.**

(A) Pathway enrichment analysis of RNA-seq data from siGATA6-transfected HPAECs showing significant upregulation of pathways related to interferon signaling, Toll-like receptor signaling, and RIG-I–like receptor signaling. The dot size represents the ratio of overlapping genes to the total number of genes in each pathway, and the color indicates the number of overlapping genes.

(B) Validation of interferon-responsive targets in independent endothelial cell lines. siRNA-mediated GATA6 knockdown in HMVEC-L. Data are means ± SD (n=5, average of five independent experiments done in triplicates). Exact p-values: *GATA6* ($p < 0.0001$), *TLR3* ($p = 0.0029$), *VCAM1* ($p = 0.0009$), *CXCL10* ($p < 0.0001$) by student t-test.

(C) Validation of interferon-responsive targets in independent endothelial cell lines. siRNA-mediated GATA6 knockdown in PromoCell-derived HPAECs significantly decreased the expression of *TLR3*, *VCAM1*, and *CXCL10*, confirming that GATA6 positively regulates these antiviral and inflammatory response genes in multiple endothelial contexts. Data are means ± SD (n=5, average of five independent experiments done in triplicates). Exact p-values: *GATA6* ($p < 0.0001$), *TLR3* ($p < 0.0001$), *VCAM1* ($p < 0.0001$) *CXCL10* ($p = 0.0001$) by student t-test.

(D) Original immunoblots corresponding to Figure 1B, showing the full, uncropped Western blot images for GATA6 (56 kDa), TLR3 (110 kDa), and β-actin (42 kDa). Boxes indicate the regions cropped and displayed in the main figure panels

**Supplementary Figure 2**: **Pyrrothiogatain treatment downregulates interferon response genes in HPAECs.**

(A) Treatment of HMVEC-L with the GATA inhibitor pyrrothiogatain (250 μM, 24 h) led to significant downregulation of *TLR3, VCAM1*, and *CXCL10* mRNA levels. Data are means ± SD (n=5, average of five independent experiments done in triplicates). Exact p-values: *GATA6* ($p < 0.0001$), *TLR3* ($p < 0.0001$), *VCAM1* ($p < 0.0001$) *CXCL10* ($p = 0.0001$) by student t-test.

(B) and PromoCell-derived HPAECs with the GATA inhibitor pyrrothiogatain (250 µM, 24 h) led to significant downregulation of *TLR3*, *VCAM1*, and *CXCL10* mRNA levels. Data are means ± SD (n=5, average of five independent experiments done in triplicates). Exact p-values: *GATA6* ($p < 0.0001$), *TLR3* ($p < 0.0001$), *VCAM1* ($p < 0.0001$) *CXCL10* ($p = 0.0001$) by student t-test.

(C) mRNA levels of *Caspase-3* and *Caspase-7* in pyrrothiogatain-treated HPAECs. Data are means ± SD (n=5, average of five independent experiments done in triplicates). Exact p-values: *CASP3* (p =ns) and *CASP7* (p =ns) by student t-test.

(D) Original immunoblots corresponding to Figure 2B, showing uncropped Western blot images for GATA6 (56 kDa), TLR3 (110 kDa), and β-actin (42 kDa). Boxed regions indicate the cropped areas displayed in the main figure panels.

**Supplementary Figure 3**: **GATA inhibition in HPAECs leads to loss of interferon gene response following PolyIC stimulation.**

(A) Human lung microvascular endothelial cells (HMVEC-L) were treated with the GATA inhibitor pyrrothiogatain (250 µM, 24 h) and/or PolyIC (10 µg/mL, 6 h) to assess GATA6-dependent induction of antiviral genes. In HMVECs, GATA6 inhibition significantly suppressed *TLR3*, *VCAM1*, and *CXCL10* mRNA induction following PolyIC stimulation. Data represents SD (n = 5; five independent experiments performed in triplicate). * $p<0.05$, **$p<0.001$, ***$p<0.0001$ by one-way ANOVA with Tukey's post hoc test.

(B) In contrast, PromoCell-derived HPAECs did not show *TLR3* induction after PolyIC treatment, while pyrrothiogatain reduced interferon-responsive genes, highlighting donor variability in PolyIC responsiveness among HPAEC lines. Data represents SD (n = 5; five independent experiments performed in triplicate). * $p<0.05$, **$p<0.001$, ***$p<0.0001$ by one-way ANOVA with Tukey's post hoc test.

(C) Original immunoblots corresponding to Figure 3E, showing uncropped Western blot images for GATA6 (56 kDa), TLR3 (110 kDa), and β-actin (42 kDa). Boxed areas denote regions cropped and presented in the main figure

**Supplementary Figure 4: Conditioned medium from GATA6 deficient HPAECs leads to upregulation**

**of interferon target genes in HPASMCs**

(A)Uncropped Western blot images corresponding to Figure 6B, showing the detection of TLR3 (110 kDa), GATA6 (56 kDa), and Tubulin (55 kDa) in PASMCs from non-diseased (control) subjects and patients with PAH.Boxed areas indicate the cropped regions displayed in the main figure.

**Supplementary Figure 5.** ***Validation of TLR3 knockdown and cell-type specificity controls**.* (A) Representative immunoblot showing efficient TLR3 silencing in HPAECs transfected with siTLR3 compared with scrambled control (siScr). β-actin was used as a loading control.

(B) Densitometry quantification of immunoblots show a >90% reduction in TLR3 protein levels following siTLR3 transfection ($p < 0.0001$, unpaired *t*-test; n = 3).

(C) Original full-length immunoblots for TLR3 (MW ≈ 110 kDa) and β-actin (MW ≈ 42 kDa) are shown, with molecular weight markers included to demonstrate antibody specificity.

(D–E) Cell identity validation by qPCR showing high endothelial marker *PECAM1* expression in pulmonary artery endothelial cells (PAECs) and high smooth muscle marker *ACTA2* expression in pulmonary artery smooth muscle cells (PASMCs), confirming cell-type purity. Data are mean ± SD; $p < 0.01$ (), $p < 0.001$ (*) by unpaired *t*-test; n = 5 biological replicates per group. Exact p-values: *PECAM1* (p =0.0010), *ACTA2* (p = 0.0015)

**Supplementary Figure 6.** ***Effect of siGATA6-conditioned media on endothelial gene expression.*** (A,B) HPAECs treated with conditioned media (CM) from siGATA6- or siScr-transfected PASMCs were analyzed for mRNA expression of *GATA6, TLR3, VCAM1, CXCL10*, and *IFIT1* by RT-qPCR. No significant differences were observed between groups, indicating that soluble factors secreted by GATA6-deficient PASMCs do not alter endothelial GATA6–TLR3–interferon signaling. Data are presented as mean ± SD (n = 5 biological replicates per group); *p* values determined by unpaired *t*-test (not significant).
